# Stomatal anatomy, leaf structure and nutrients of tropical rainforest tree species respond to altitude in a coordinated manner in accordance with the leaf economics spectrum

**DOI:** 10.1101/2022.01.16.476499

**Authors:** Bhagya Weththasinghe, Iroja Caldera, Nimalka Sanjeewani, Dilum Samarasinghe, Himesh Jayasinghe, Asanga Wijethunga, Janendra De Costa

## Abstract

Understanding taxon level responses of key plant traits to environmental variation across tropical rainforests (TRFs) is important to determine their response to climate change. We used an altitudinal gradient (from 150 to 2100 m above sea level, asl) across TRFs in Sri Lanka to answer the following questions: (a) Does the response to altitude by stomatal traits differ among plant taxa in TRFs?; (b) Are the altitudinal responses of key leaf structural traits (e.g. leaf mass per area, LMA) and major leaf nutrient (nitrogen, N, and phosphorus, P) concentrations linked to the corresponding variation in stomatal traits in a coordinated response across taxa?; (c) How strong is the influence of climatic variation on responses of leaf traits to altitude?. Leaf samples were collected in permanent sampling plots within rainforest reserves at Kanneliya (150 m asl), Sinharaja-Enasalwatta (1050 m), Hakgala (1800 m) and Piduruthalagala (2100 m) from 19 species in three plant genera (*Calophyllum*, *Semecarpus* and *Syzygium*). Stomatal density, guard cell length and epidermal density showed variation among taxa, but did not respond to altitude. Potential conductance index (PCI), a proxy for photosynthetic capacity, decreased with increasing altitude, in a common response across taxa. We found evidence that altitudinal responses of LMA, leaf N and P were linked to stomatal responses in a coordinated manner, where key features were the negative correlations between PCI and LMA and between proxy photosynthetic N- and P-use efficiencies (‘PNUE’ and ‘PPUE’) and LMA. We found strong responses to climatic variation across taxa and altitudes, where PCI, ‘PNUE’ and ‘PPUE’ increased and LMA decreased with increasing temperature, precipitation and solar irradiance. We conclude that stomatal traits of tree species in TRFs form part of a coordinated leaf trait response to environmental change which is in accordance with the leaf economics spectrum.

## Introduction

Tropical rainforests (TRFs) are one of the most sensitive ecosystems to environmental change (Malhi et al. 2009, 2014, Zelazowski et al. 2011, Hubau et al. 2020). The high diversity of plant species in TRFs are likely to elicit different responses by different species to changes in climatic and soil variables (Hofhansl et al. 2020). Therefore, it is important to determine the species/genus level variation of responses to understand and predict ecosystem responses to environmental perturbations such as long-term climate change and climate extremes.

Altitudinal gradients are used as a means of detecting the influence of environmental change on tropical ecosystems, their processes and productivity (Malhi et al. 2010, 2017). Ecosystems such as TRFs which span a wide range of altitudes experience changes in several key environmental variables that vary with altitude. For example, air and soil temperatures, atmospheric pressure, vapour pressure deficit and partial pressure of CO_2_ (pCO_2_) decrease with altitude whereas solar irradiance, wind speed and incident UV radiation could increase (Barry 1992, Körner 2007). Such changes in climatic variables along altitudinal gradients have induced altitude-linked variations in soil properties by influencing key soil processes such as mineralization and organic matter decomposition (Dieleman et al. 2013). Evolutionary and adaptive responses of plants to altitude-linked gradients in climate and soil variables have led to variations in species composition and diversity in TRFs along altitudinal gradients (Lieberman et al. 1996, Vazquez G and Givnish 1998, Aiba and Kitayama 1999, Givnish 1999). Furthermore, differences in key plant functional traits/leaf traits have been observed across plant species that inhabit altitudinal gradients (Oliveras et al. 2020).

Stomata occupy a strategic location in plants to regulate their carbon and water balances via leaf-air exchange of CO_2_ and water vapour. Therefore, responses of stomatal density and size to altitude represent a key pathway to optimize carbon and water balances in the face of environmental variation across an altitudinal gradient. Many early work reported increased stomatal density (SD) and stomatal index (SI) with increasing altitude (Korner and Cochrane 1985, Korner et al. 1986, McElwain 2004, Kouwenberg et al. 2007), which was interpreted as confirmation of the decrease of SD and SI with increasing pCO_2_, reported by (Woodward 1987, Woodward et al. 2002). However, a broader survey of literature shows a diverse range of stomatal responses to altitude. Accordingly, increased SD and SI with increasing pCO_2_ along altitudinal gradients (Hu et al. 2015) and time (Bai et al. 2015) have been reported. Furthermore, increases in SD and SI up to 2800 – 3000 m above mean sea level (asl) have been observed to reverse in some plant species at higher altitudes (Qiang et al. 2003, Li et al. 2006, Luo et al. 2006). In comparison to SD and SI, response of stomatal size, measured as the product between guard cell length (GCL) and closed stomatal width, to environmental variation has been studied less frequently (Lomax et al. 2009). A highly-conserved inverse relationship has been shown between stomatal density and size across different taxonomic groups and environmental variables (Franks and Beerling 2009, Doheny-Adams et al. 2012). However, GCL of *Arabidopsis thaliana* ecotypes from a range of altitudes from 50 to 1260 m amsl showed all possible responses (i.e. increase, decrease and no change) to increased pCO_2_ while not showing significant correlations with either SD or SI (Caldera et al. 2017). The above range of stomatal responses illustrates the inter- and intra-specific diversity in this response. Accordingly, in an ecosystem of high species diversity such as the TRFs, a diversity of stomatal responses to altitude could be expected. Long-term evolutionary responses and short-term acclimation to environmental variation across altitudes probably combine to bring about these diverse stomatal responses (Hultine and Marshall 2000, Qiang et al. 2003, Franks and Beerling 2009, Haworth et al. 2015).

As stomatal anatomy influences the assimilate supply for construction of leaves, altitudinal variation of leaf structural traits such as the leaf mass per area (LMA) could be linked to the corresponding variation of stomatal traits. Leaf mass per area is a measure of assimilate investment per unit area of photosynthetic surface and is regarded as a key trait of the leaf economic spectrum (Wright et al. 2004). Shi et al. (2015) showed that diverse responses of stomatal traits to altitude were linked to leaf economic strategy and plant growth habit. Nitrogen (N) and phosphorus (P) are the two foremost plant nutrients and are considered most limiting for plant growth (Reich and Oleksyn 2004). Nitrogen is an essential component of enzymes, especially the key photosynthetic enzyme Rubisco, while P is a key part of nucleic acids, cell membranes and energy-carriers such as ATP and NADPH. Leaf N and P concentrations are indicative of their availability in the soil as well as their proportional allocation to leaves to meet the physiological demands of processes that require N (e.g. enzymes) and P (e.g. ATP and nucleic acids). Accordingly, variation in leaf N and P concentrations across an altitudinal gradient, when considered along with corresponding variations in stomatal and leaf structural traits could be part of an integrated ecosystem level response to environmental variation as represented by altitude (Oliveras et al. 2020). The role of genetic variation, based on taxonomic grouping, as opposed to variation caused by local environmental factors is poorly understood for leaf traits in different plant species in complex ecosystems such TRFs.

Tropical rainforests of Sri Lanka (TRFSL) are located in the humid tropical climatic zone of Sri Lanka in its south-western plains and the western slope of its central highlands. Lowland rainforests are found in the lower altitudes up to ca. 1000 m amsl while montane forests are found in the higher altitudes up to 2200 m. Species richness and diversity of TRFSL have been shown to decrease with increasing altitude (Sanjeewani et al. 2020) and there are no plant species inhabiting the whole altitudinal range of the TRFSL (Sanjeewani et al. Unpublished). Therefore, genetically-determined variations in stomatal and leaf traits among different species inhabiting different altitudes would confound responses of these traits to environmental gradients across altitudes. Furthermore, proportional contributions from genetic and environmental components to the observed variation of these traits could be different for different species (Zhang et al. 2012). In this study, we examine altitudinal variation of stomatal traits in three plant genera that have different species inhabiting a sufficiently wide altitudinal gradient with the objective of identifying variation patterns of their stomatal traits with altitude and then elucidating their possible environmental and genus-level genetic controls. As a second objective, we examine possible inter-relationships between stomatal and leaf traits with a view to determine possible trait assemblies that underpin the response of these traits to specific climatic variation across altitudes. We asked the following questions: (a) Does the response to altitude by stomatal anatomical traits differ among plant taxa (e.g. genera) in TRFs?; (b) Are the altitudinal responses of key leaf structural traits (LMA) and major leaf nutrient (N and P) concentrations linked to the corresponding variation in stomatal traits in a coordinated response across different taxa (e.g. genera, species)?; (c) How strong is the influence of climate variation across altitudinal gradients on responses of leaf traits to altitude?. Specifically, we tested the following hypotheses: Hypothesis 1: There is significant variation among taxa in the response of their stomatal traits to variation in altitude; Hypothesis 2: Altitudinal responses of key leaf structural traits, major leaf nutrient concentrations and stomatal traits form a coordinated response across different taxa in tropical rainforests; Hypothesis 3: Responses of leaf traits to altitude are strongly influenced by climatic variables that vary across altitudinal gradients.

## Materials and methods

### Study sites and climate

Collection of leaf samples for the study was carried out in six permanent sampling plots (PSPs) of 1 ha (100 m x 100 m) each established along an altitudinal gradient from 117 m to 2080 m asl (Fig. 1). All plots were located in the undisturbed areas inside selected TRFSL. Two plots were located in the Kanneliya Forest Reserve. Another two plots were located in the Sinharaja Forest Reserve (Enasalwatte) which is a World Heritage site. The other two plots were located in the Hakgala Strict Nature Reserve and Piduruthalagala Forest Reserve. Long-term (1970-2000) averages of the climate variables of study sites were obtained from the high-resolution (1 km^2^) global climatic database WorldClim 2 (Fick and Hijmans 2017) (Table 1). Long-term annual averages of the mean, maximum and minimum air temperatures, vapour pressure, wind speed and daily solar irradiance were computed from their monthly means. Long-term average annual precipitation was computed from the respective annual totals.

**Table 1.** Study sites and their long-term (1970-2000) climatic variables.

| Location | Latitude<br>(N) | Longitude<br>(E) | Altitude<br>(m) | T<sub>AV</sub><br>(°C) | T<sub>max</sub><br>(°C) | T<sub>min</sub><br>(°C) | R<sub>F</sub><br>(mm y<sup>-1</sup>) | Radiation<br>(kJ m<sup>-2</sup> d<sup>-1</sup>) | WVP<br>(kPa) | WS<br>(ms<sup>-1</sup>) |
| --- | --- | --- | --- | --- | --- | --- | --- | --- | --- | --- |
| KDN 1 | 6.24749 N | 80.34071 E | 117 | 26.3 | 29.8 | 23.5 | 3656.0 | 18945.3 | 2.8 | 2.3 |
| KDN 2 | 6.26045 N | 80.35112 E | 174 | 26.3 | 29.3 | 22.8 | 3788.0 | 18832.8 | 2.8 | 2.3 |
| ENS 2 | 6.39433 N | 80.59709 E | 1042 | 21.3 | 24.5 | 18.1 | 3103.0 | 17910.3 | 2.0 | 2.2 |
| ENS 2 | 6.39441 N | 80.59652 E | 1065 | 21.3 | 24.5 | 18.1 | 3103.0 | 17910.3 | 2.0 | 2.2 |
| HKG | 6.92725 N | 80.81839 E | 1804 | 17.2 | 20.3 | 13.8 | 2017.0 | 17688.3 | 1.8 | 2.0 |
| PTG | 6.98197 N | 80.77276 E | 2080 | 15.3 | 18.4 | 12.3 | 1903.0 | 17460.0 | 1.2 | 1.9 |

KDN 1 & 2 – Kanneliya Forest Reserve Plots 1 & 2; ENS 1 & 2 – Sinharaja-Enasalwatte Plots 1 & 2; HKG – Hakgala Strict Nature Reserve; PTG – Pidurutalagala Forest Reserve. T<sub>AV</sub> – Mean annual average temperature; T<sub>max</sub> - Mean annual maximum temperature; T<sub>min</sub> - Mean annual minimum temperature; R<sub>F</sub> - Mean annual total precipitation; Radiation – Mean daily incident solar radiation; WVP - Mean atmospheric vapour pressure; WS – Mean wind speed (Source: Fick and Hijmans, 2017).

**Figure 1.**
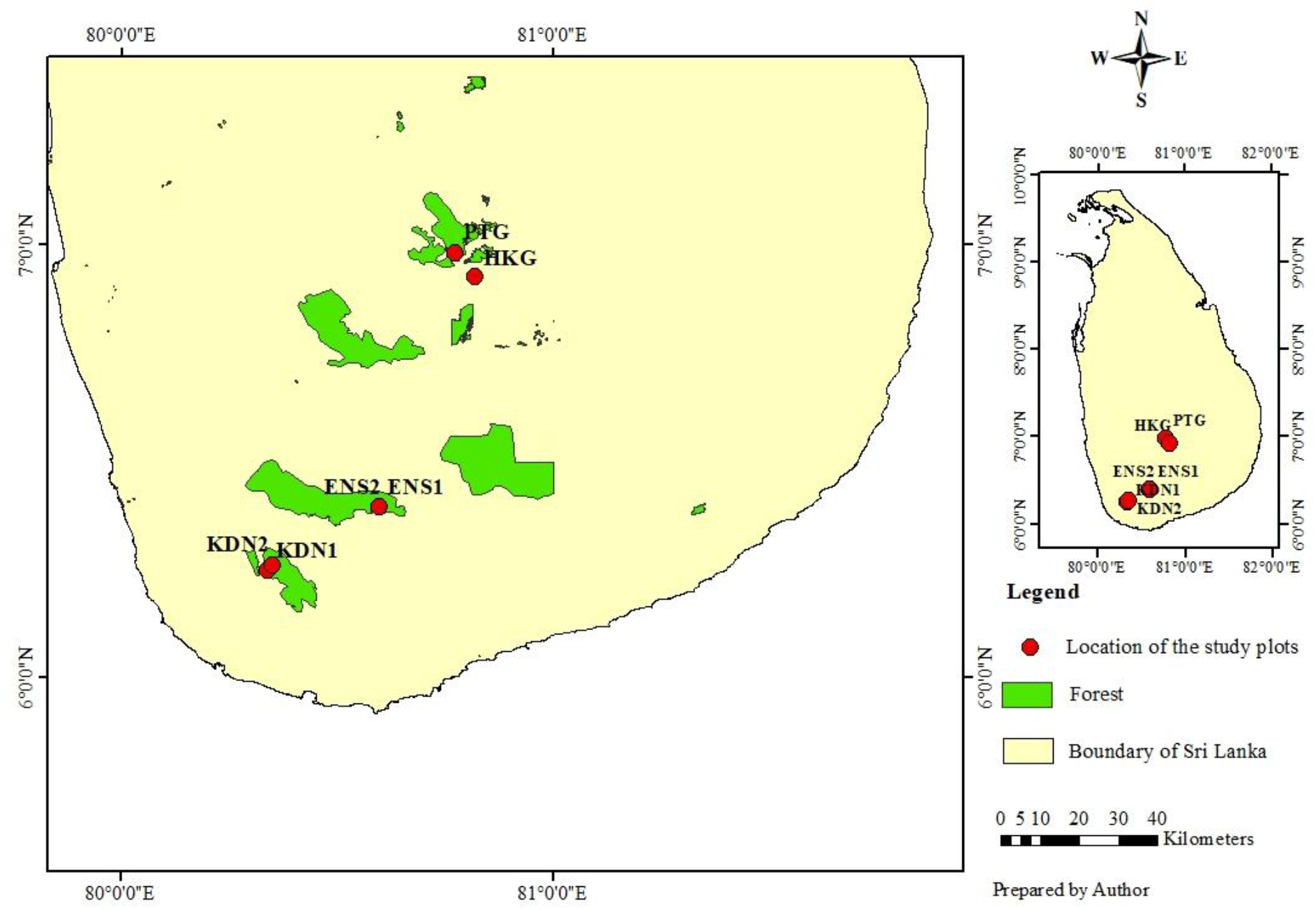
Permanent sampling plots of the present study located at different altitudes in tropical rainforests of Sri Lanka. Kanneliya Plot 1 (KDN1-117 m asl), Kanneliya Plot 2 (KDN2-174 m asl), Sinharaja-Enasalwatta Plot 1 (1042 m asl), Sinharaja-Enasalwatta Plot 2 (1065 m asl), Hakgala Plot (1804 m asl) and Piduruthalagala Plot (2080 m asl).

### Plant species

A complete census of all plants having a diameter at breast height (1.3 m above ground level or buttress) of 10 cm or greater had been carried out in the PSPs in 2019 (Sanjeewani et al. 2020). Plant species for the study were selected based on the availability of a species across a sufficiently wide altitudinal range. However, except *Semecarpus gardneri*, which was present in plots at 150 m and 1050 m, there was no other species inhabiting more than one altitude. At the genus level, there were three genera, viz. *Syzygium*, *Semecarpus* and *Calophyllum* which had different species occupying a wide altitudinal range. Accordingly, 19 species from these genera were selected for the study (Table 2).

**Table 2.** Species from selected genera for the study and their locations.

| Location | Altitude (m) | *Syzygium* | *Semecarpus* | *Calophyllum* |
| --- | --- | --- | --- | --- |
| Kanneliya | 150<sup>†</sup> | *S. alubo*<br>*S. neesianum*<br>*S. firmum* | *S. walkeri*<br>*S. gardneri*<br>*S. subpeltata* | *C. bracteatum*<br>*C. cordato-oblongum* |
| Sinharaja-Enasalwatte | 1050<sup>†</sup> | *S. cylindricum*<br>*S. micranthum*<br>*Syzygium* new sp. | *S. gardneri*<br>*S. parvifolia* | *C. acidus* |
| Hakgala | 1800 | *S. paniculatum*<br>*S. zeylanicum* | *S. obovata* |  |
| Pidurutalagala | 2100 | *S. revolutum*<br>*S. rotundifolium* |  | *C. walkeri* |

<sup>†</sup>Altitude at mid-point between the two plots at Kanneliya and Sinharaja-Enasalwatte was selected.

The three core-eudicot genera belong to the clade Rosids and are classified under three different orders, viz., *Calophyllum* (Calophyllaceae) in Malpighiales, *Semecarpus* (Anacardiaceae) in Sapindales and *Syzygium* (Myrtaceae) in Myrtales (Angiosperm Phylogeny Group 2009). *Calophyllum* is a pantropical genus consisting of approximately 187 species. In Sri Lanka, 11 out of 13 *Calophyllum* species are endemic and are restricted to rainforests (Kostermans 1980). About 50 species of *Semecarpus* species are distributed from tropical Asia (Indo-Malaysian region) to Oceania. All 12 species of *Semecarpus* in Sri Lanka are endemic and mainly distributed in low- to mid-altitude rainforests (Meijer 1983). Approximately 1200 species of *Syzygium* are found in tropical Africa, subtropical to tropical Asia, Australia, New Caledonia, New Zealand and Pacific islands. There are 32 *Syzygium* species among Sri Lankan flora and 25 of them are endemic. The genus has a wide altitudinal range in Sri Lanka with a majority of species in higher-altitude rainforests. There are some introduced species, usually under cultivation while some are naturalized (Ashton 1981).

### Sample collection

Samples of mature, healthy leaves were taken from harvested branches of trees of the above 19 species. Three leaves per plant from five plants per species were used for stomatal measurements. Species from two PSPs at near equal altitudes (Kanneliya and Enasalwatta) were pooled for measurements and data analysis. All measurements were done on fully-expanded mature leaves of each branch.

### Measurement of stomatal traits

Using high precision Polyvinylsiloxane (Muller-Omicron GmbH & Co. KG, Germany) negative impressions were acquired from leaves within 24 hours following harvesting of branches, before leaf wilting occurred. Impressions were taken from a leaf position in the middle between mid-rib and leaf margin and in the middle between the apex and the base of the leaf. A preliminary study on each sampled species revealed that stomata are present only on the abaxial leaf surface. Therefore, the negative impressions were taken only from the abaxial leaf surface.

Afterwards transparent cellulose varnish was applied on the negative silicone rubber impressions. The varnish was allowed to dry off and then the positive impressions of the leaf abaxial surfaces were peeled off. Thereafter, they were mounted onto microscopic slides and covered with coverslips which were secured with adhesive tape. Five random fields per leaf sample were selected and the images of these fields were taken under light microscopy (Primo Star, Zeiss, Germany). This was done by a camera coupled with the microscope and Zeiss Zen microscope software. Using these microscopic images (Plates S1-S5), number of stomata and epidermal cells per microscopic field were counted.

All sampled species are rainforest tree species. Therefore, the borders of epidermal cells were difficult to differentiate in the leaf imprints. To overcome this difficulty all images were sharpened and the black and white contrast was adjusted using Adobe Photoshop 7.0.1. The multi-point tool of the ImageJ software was used when counting stomata and epidermal cells. Editing of images and use of the multi-point tool enabled precise counts of stomata and epidermal cells.

Stomatal density (SD) and epidermal density (ED) were calculated as the number of stomata and epidermal cells per unit leaf area (mm^-2^). Stomatal index (SI) was computed as the ratio between stomatal number and the number of total epidermal cells, including stomatal guard cells, using the formula, SI = (SD/[SD + ED] x 100).

Guard cell length (GCL) was measured using the slides prepared for determination of stomatal traits. Measurements were taken by the Line tool of the Graphics in UI Automation mode of the Zeiss Zen Microscope Software. This was done for one stomate each from randomly selected five fields of view per leaf sample.

Potential conductance index (PCI) was computed as PCI = [GCL] ^2^ x SD x 10^-4^ (Holland and Richardson 2009), where GCL is given in μm and SD in stomata per mm^2^, assuming that the stomatal aperture area is proportional to the GCL^2^. As PCI combines measures of maximum stomatal size (GCL) with stomatal density (SD), it can be considered a measure of maximum stomatal conductance (g_max_) (Franks and Beerling 2009).

### Measurement of leaf structural traits and nutrients

Leaf blade area (LBA) was measured on fully-expanded leaves by tracing the outline of the leaves on a white paper. A printed scale was kept on the white paper and images were taken by a camera. Leaf blade area was measured using these images using ImageJ software. The leaf samples were dried to constant weight at 105°C in an oven (MOU-1125, SANYO, Japan). Each leaf was weighed separately using an electronic balance (AR2130, Obaus Co., USA). Leaf mass per area was computed as the ratio between leaf dry mass and LBA. Taking PCI as a proxy for photosynthetic capacity (Wong et al. 1979, Farquhar and Sharkey 1982), proxy indices of photosynthetic nitrogen and phosphorus use efficiencies (‘PNUE’ and ‘PPUE’ respectively) were computed as the respective ratios of PCI with leaf N and P.

### Determination of N and P concentrations of leaf tissues

Leaf N concentration was measured using the modified Kjeldahl method (Motsara and Roy 2008). Leaf P concentration was measured by dry ashing at 450°C followed by digestion in HNO_3_. Phosphorus concentration of the ash extract was determined by the spectrophotometric vanadium phosphomolybdate method (Motsara and Roy 2008).

### Statistical analysis

An initial exploration of the distributions of measured variables within each species at each altitude was done with box plots. Normality of distributions was tested by the Shapiro-Wilk statistic computed with Proc Univariate in SAS^®^ Studio (Version 9.4). The influence of genus and altitude on the measured traits was examined by a linear mixed model (LMM). As the three genera have been selected from among a large number of genera in the PSPs, the genus effect was considered as random. The altitude effect was considered as fixed because the selection of altitudes for leaf sampling was constrained to those where PSPs have been established. This LMM analysis was done for each trait using Proc Mixed in SAS with the residual (restricted) maximum likelihood method. The genus x altitude interaction was considered a random effect.

Inter-relationships among stomatal anatomical traits were determined by linear correlation analysis using all replicate measurements (i.e. n=1425). In order to examine inter-relationships among the leaf anatomical and structural traits and leaf nutrients, linear correlation analysis was done on the means of all leaf traits of the 19 plant species at different altitudes. The respective variables were averaged across altitudes, replicate trees, leaves and microscope fields when calculating species means. Influence of individual climatic variables on measured leaf traits were determined by simple linear regression. Data from all species and altitudes were pooled in the regressions between leaf traits and climatic variables. *Semecarpus gardneri* was the only species which was present at more than one altitude (i.e. at 150 and 1050 m). Mean values from *S. gardneri* at the two altitudes were considered as two separate data points in the correlation and regression analyses.

A factor analysis (Child 1990) was done to identify underlying factors that cause variation in leaf traits in the 19 plant species belonging to the three genera that are distributed across the altitudinal gradient. In this analysis also, *S. gardneri* at 150 and 1050 m were considered as two observations. The factor analysis was done using the principal axis method with prior communality estimates as one. All factors that showed Eigen values greater than one were extracted and varimax rotation was performed to identify variables that loaded on to extracted factors. Factor scores of the 20 observations were plotted in the factor space of the two principal factors (i.e. the two factors having the highest Eigen values) to determine aggregation/separation of the 19 species based on their leaf traits. A cluster analysis using the complete linkage method was performed to determine species clusters among the 19 species, based on their leaf traits. Multivariate analysis of variance (MANOVA) and the Wilks’ Lambda Statistic was used to determine the significance of differences between clusters. All statistical analyses were performed using SAS^®^ Studio (Version 9.4).

## Results

### Distributions of measured stomatal traits at different altitudes

All stomatal traits showed substantial variation among species at a given altitude (Fig. S1). Species-wise distributions of most stomatal traits showed normality in a majority of species whereas most distributions of stomatal index deviated from normality (Table S1). The few outliers were located on both ends of the distributions.

### Influence of genus and altitude on stomatal traits

Analysis of stomatal traits in a linear mixed model showed that the fixed effect of altitude was significant (p=0.06) on potential conductance index (PCI), but was non-significant (p>0.07) on all other stomatal traits (Table 3). The random effect of the genus contributed a higher percentage (56-64%) of the total variance of guard cell length (GCL), stomatal density (SD) and epidermal density (ED) in comparison to the genus x altitude interaction (10-26%) (Table 4). The genus effect contributed very little to the total variance of stomatal index (SI) and PCI (0-4%).

**Table 3.** Results of Type III tests for fixed altitude effect on leaf traits of plant species in tropical rainforests of Sri Lanka along an altitudinal gradient.

|  | GCL | SD | ED | SI | PCI | LMA | LBA | N | P | N:P |
| --- | --- | --- | --- | --- | --- | --- | --- | --- | --- | --- |
| $F_{3,4}$ | 1.21 | 0.50 | 1.47 | 1.65 | 5.84 | 26.30 | 17.87 | 13.72 | 2.33 | 0.73 |
| $p>F$ | 0.414 | 0.702 | 0.350 | 0.313 | 0.061 | 0.0043 | 0.0088 | 0.0143 | 0.2161 | 0.5840 |

GCL – Guard cell length; SD – Stomatal density; ED – Epidermal density; SI – Stomatal index; PCI – Potential conductance index; LMA – Leaf mass per unit area; LBA – Leaf blade area; N – Leaf nitrogen concentration; P – Leaf phosphorus concentration; N:P – Leaf nitrogen: phosphorus ratio.

### Variation of stomatal traits of different genera with altitude

Guard cell length of *Calophyllum* was greater than that of *Semecarpus* and *Syzygium* at all altitudes (Fig. 2.a). There was no consistent trend with altitude for GCL of any genus, thus confirming the absence of an altitude effect (Table 3). The two notable increases in GCL with increasing altitude in *Calophyllum* at 1050 m and *Semecarpus* at 1800 m occurred because of a single species (*Calophyllum acidus* and *Semecarpus obovata*, Fig. S1.a) having a substantially higher GCL than others within the respective genera.

**Figure 2.**
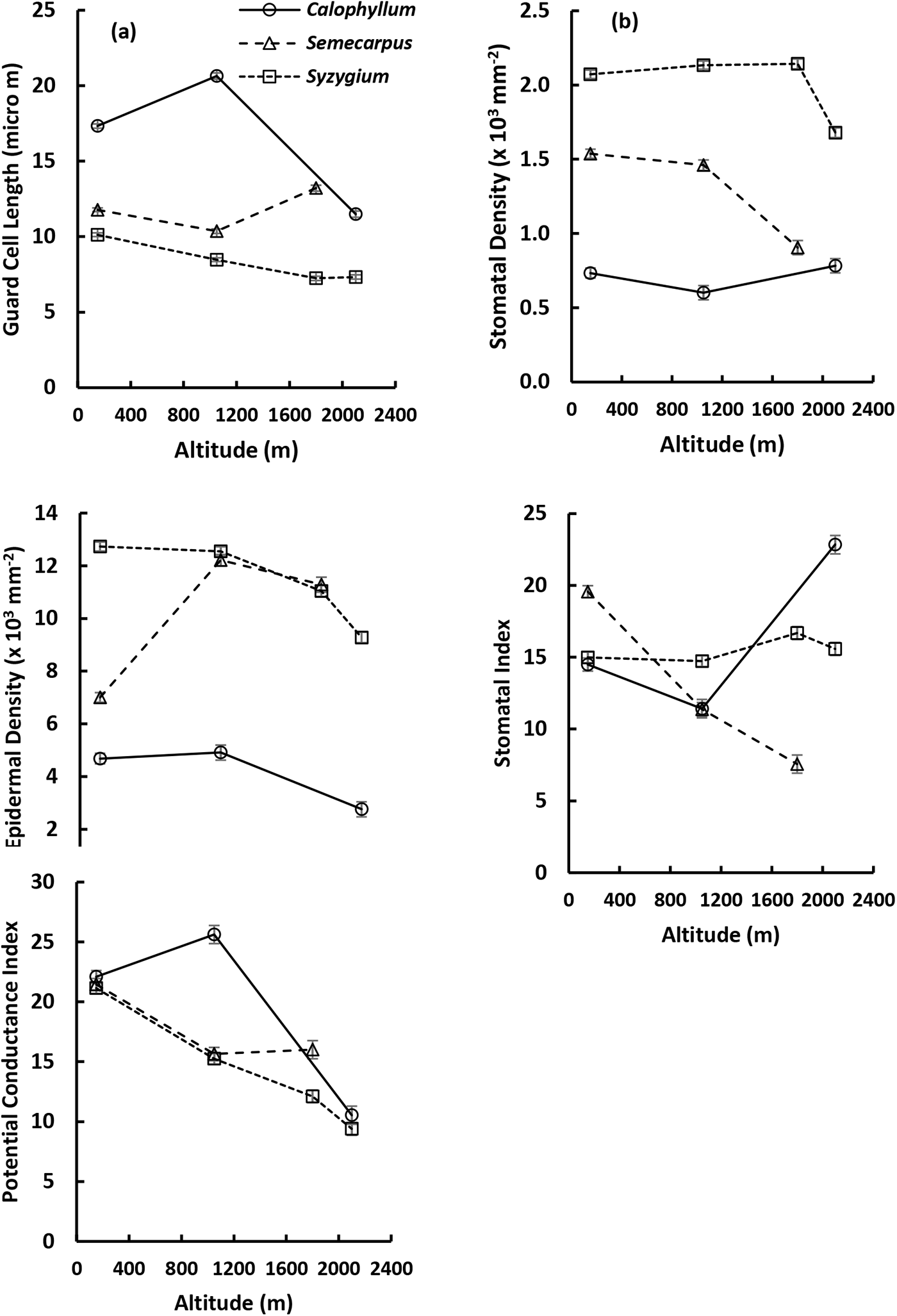
Variation of least square (LS) means of stomatal traits of different plant genera with altitude: (a) Guard cell length; (b) Stomatal density; (c) Epidermal density; (d) Stomatal index; (e) Potential conductance index. Each data point is the LS mean of different species of each genus at a given altitude. Error bars indicate standard errors of LS means. Number of data points for each LS mean varied from 75 to 225.

Stomatal density (SD) showed clear separation between the three genera with *Syzygium* having the highest and *Calophyllum* the lowest with *Semecarpus* intermediate (Fig. 2.b). In all genera, variations of SD with altitude were smaller than the variation among genera, thus confirming the higher variance component attributed to genus (Table 4). The two notable reductions of SD at higher altitudes in *Syzygium* and *Semecarpus* had occurred because of species level variation (*Syzygium rotundifolium*, *Syzygium revolutum* and *Semecarpus obovata*) (Fig. S1.b). These reductions contributed to the genus x altitude variance component. There is a highly-significant (*p*<0.0001) negative correlation between GCL and SD (*r* = -0.67; n = 1425) (Table S2). Similar correlations were evident at the genus level as well (*Syzygium*: *r* = -0.160, p<0.0001, n = 735; *Calophyllum*: *r* = -0.312, p<0.0001, n = 300; *Semecarpus*: *r*=-0.335, p<0.0001; n = 390).

Epidermal density (ED) of *Calophyllum* was substantially lower than those of *Semecarpus* and *Syzygium* (Fig. 2.c). Even though ED of all genera showed a decreasing trend from 1050 m onwards, the observed reductions were smaller than differences among genera, which confirmed the greater variance component due to the genus effect (Table 3). The lower ED of *Semecarpus* at 150 m, which occurred because of lower ED of all three *Semecarpus* species (i.e. *S*. *subpeltata*, *S. walkeri* and *S. gardneri*) at this altitude (Table 2), contributed to genus x altitude variance (Table 4). Epidermal density showed a positive correlation with SD and a negative correlation with GCL (Table S2).

Stomatal index (SI) did not show a consistent variation with altitude or genus (Fig. 2.d). Accordingly, the variance component due to genus was zero (Table 4). Furthermore, the variance component attributed to genus x altitude interaction (32%) was lower than the residual (68%). Stomatal index was negatively-correlated with ED and GCL and positively-correlated with SD (Table S2).

**Table 4.** Estimates of variance components for random effects on stomatal traits in plant species of tropical rainforests in Sri Lanka across an altitudinal gradient.

| Effect | GCL | % | SD | % | ED | % | SI | % | PCI | % |
| --- | --- | --- | --- | --- | --- | --- | --- | --- | --- | --- |
| Genus | 16.517 | 63 | 428810 | 64 | 12190137 | 56 | 0 | 0 | 2.341 | 4 |
| Genus x Altitude | 6.678 | 26 | 68110 | 10 | 3178939 | 15 | 14.875 | 32 | 10.053 | 18 |
| Residual | 2.982 | 11 | 169329 | 25 | 6212000 | 29 | 31.107 | 68 | 42.550 | 77 |
| ML | 5646.7 |  | 21193.9 |  | 26313.4 |  | 8965.3 |  | 9407.4 |  |
| AIC | 5652.7 |  | 21199.9 |  | 26319.4 |  | 8969.3 |  | 9413.4 |  |
| BIC | 5650 |  | 21197.2 |  | 26316.7 |  | 8967.5 |  | 9410.6 |  |

ML: -2Res Maximum Likelihood statistic; AIC: Akaike Information Criterion; BIC: Bayesian Information Criterion.

Potential conductance index (PCI) showed a general decreasing trend with altitude (Fig. 2.e) which confirmed the significance of the fixed effect of altitude (Table 3). Furthermore, PCI did not show variance due to genus. The high PCI of *Calophyllum* at 1050 m, which was an exception to the decreasing trend with altitude, occurred because of the high GCL of *C. acidus* (Fig. S1.a). Potential conductance index was negatively-correlated with GCL (Table S2). PCI also showed a positive correlation with SI and a negative correlation with ED, but both were very weak (*r* < 0.12).

### Variation of leaf structural traits of different genera with altitude

Distributions of leaf mass per area (LMA) and leaf blade area (LBA) showed variation among different species at each altitude (Fig. S2). A large majority of the species x altitude combinations showed normal distributions in their LMA and LBA (Table S1) with very few outliers.

The fixed effect of altitude on LMA and LBA were highly-significant (Table 3). The random effect of the genus did not contribute to variance of LMA (Table 5). In contrast, the variance due to genus contributed 46% to the total variance of LBA. Leaf mass per area showed increasing trends with altitude in all three genera (Fig. 3.a). In contrast, LBA of all genera showed decreasing trends (Fig. 3.b). There was clear separation between genera with *Semecarpus* having greater LBA than the other two.

**Figure 3.**
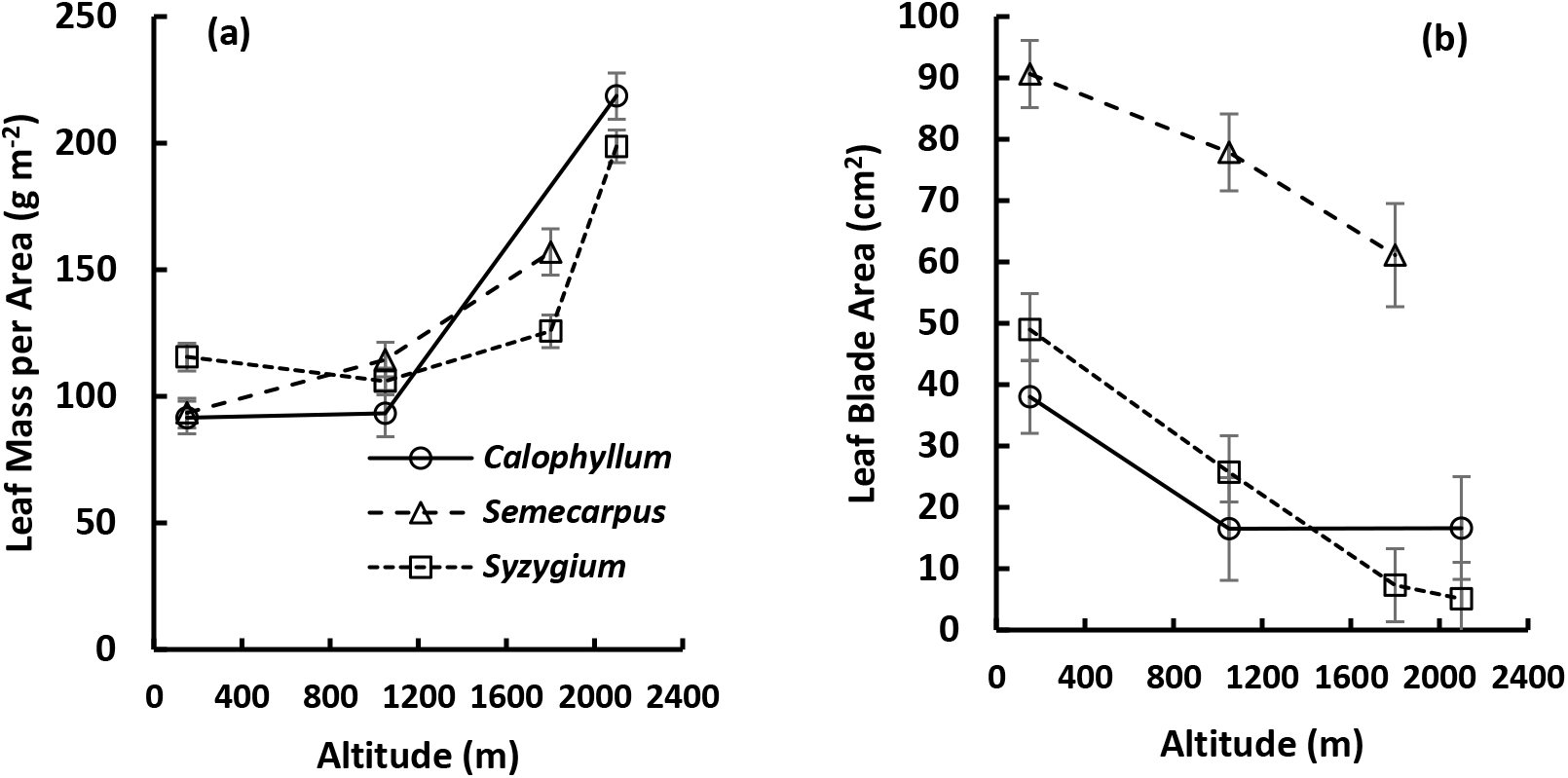
Variation of LS means of leaf structural properties of different plant genera with altitude: (a) Leaf mass per area; (b) Leaf blade area. Number of data points for each LS mean varied from 15 to 45.

**Table 5.** Estimates of variance components for random effects on leaf structural traits and nutrients in plant species of tropical rainforests in Sri Lanka along an altitudinal gradient.

| Effect | LMA | % | LBA | % | N | % | P | % | N:P | % |
| --- | --- | --- | --- | --- | --- | --- | --- | --- | --- | --- |
| Genus | 0 | 0 | 892.6 | 45.9 | 0 | 0 | 8333 | 23 | 53.7 | 28 |
| Genus x Altitude | 156.9 | 11 | 1.44 | 0.1 | 0.574 | 14 | 0 | 0 | 0 | 0 |
| Residual | 1242.5 | 89 | 1049.9 | 54.0 | 3.576 | 86 | 27304 | 77 | 140.9 | 72 |
| ML | 2825.1 |  | 2777.6 |  | 244.6 |  | 731.9 |  | 434.8 |  |
| AIC | 2829.1 |  | 2783.6 |  | 248.6 |  | 735.9 |  | 438.8 |  |
| BIC | 2827.3 |  | 2780.9 |  | 246.8 |  | 734.1 |  | 437.0 |  |

ML: -2Res Maximum Likelihood statistic; AIC: Akaike Information Criterion; BIC: Bayesian Information Criterion.

### Variation of leaf nutrients of different genera with altitude

Distributions of leaf nitrogen (N) and phosphorus (P) concentrations and their ratio (N:P) showed variation among species and altitudes, especially in their spread (Fig. S3). However, all but one (N) or two (N:P) species showed normality in their distributions (Table S1). Because of the lower number of data points at the species level, distributions of leaf nutrients were examined at the genus level as well. The distributions of leaf nutrients of different genera showed a narrower spread, especially for P and N:P (Fig. S4). All but one (N:P) and two (P) distributions were normal (Table S3).

The fixed effect of altitude was significant (p<0.05) for leaf N, but non-significant for P and N:P (Table 3). The random effect of genus did not contribute to the variance of leaf N, but contributed 23% and 28% respectively to the variances of P and N:P (Table 5). The random effect of genus x altitude did not contribute to the variances of P or N:P while contributing 14% to that of N.

All three genera showed increased leaf N from 150 m to 1050 m (Fig. 4), which was followed by decreases in *Calophyllum* and *Semecarpus* at higher altitudes. In *Syzygium*, leaf N continued to increase up to 1800 m, which was followed by a steep decline at 2100 m. Leaf P and N:P did not show consistent trends with altitude. Differences in leaf nutrients among genera were not consistent across altitudes. *Syzygium* had higher leaf N at 1800 m whereas *Semecarpus* had lower leaf P had 150 m. *Calophyllum* had higher leaf N:P at 150 m and 2100 m.

**Figure 4.**
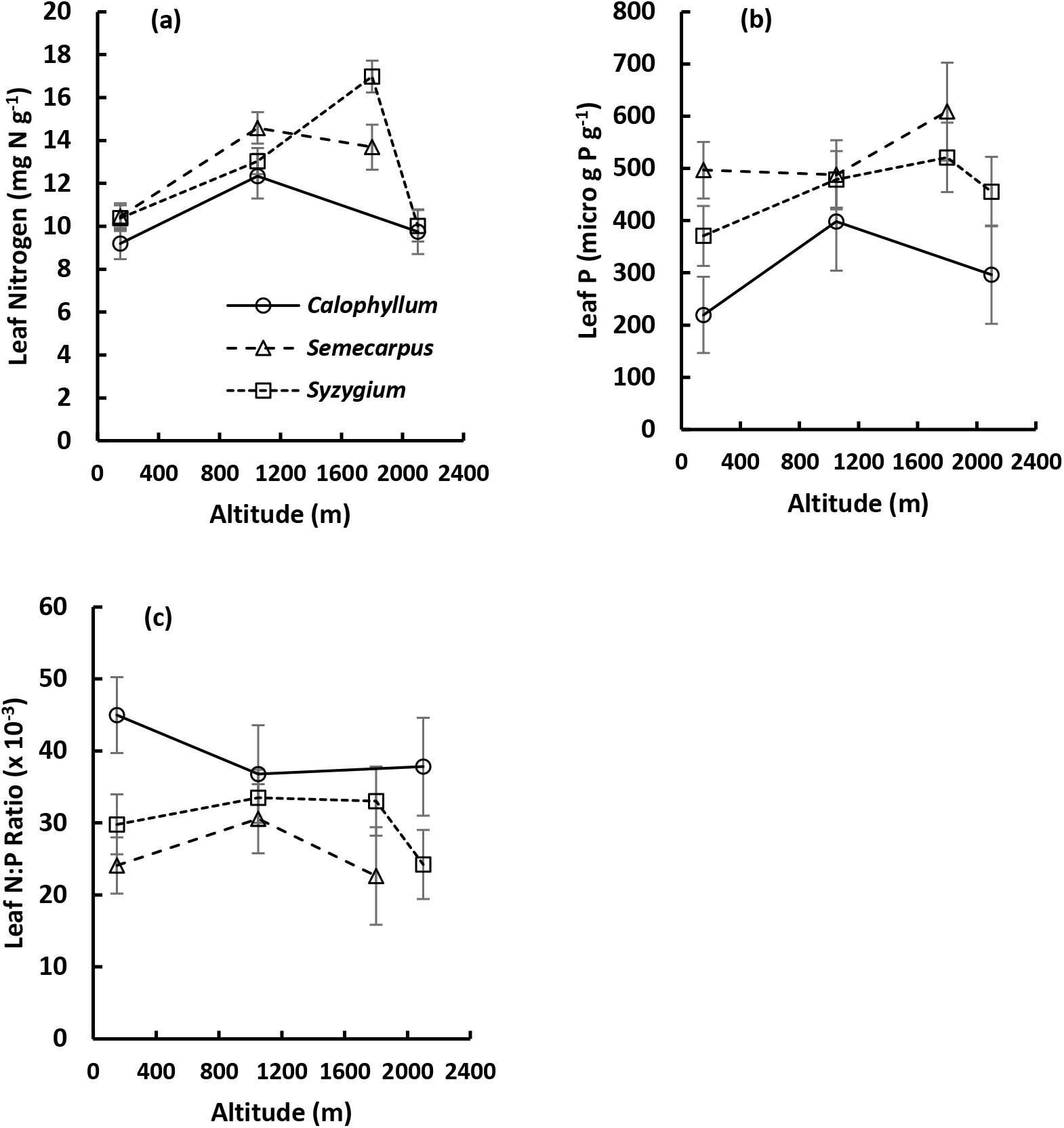
Variation of LS means of leaf nutrients of different plant genera with altitude: (a) Leaf N; (b) Leaf P; (c) Leaf N:P ratio. Error bars indicate standard errors of LS means. Number of data points for each LS mean varied from 3 to 9.

### Inter-relationships among measured variables

Table 6 shows the linear correlation matrix among species means of leaf stomatal and structural traits and leaf nutrients. Guard cell length (GCL) showed significant (p<0.05) negative correlations with SD and ED, but had a positive correlation with PCI. Stomatal density (SD) was positively correlated with ED, which in turn was negatively correlated with SI. Except for a positive correlation between ED and leaf N, there were no significant correlations between stomatal anatomical traits and leaf nutrients.

**Table 6.** Linear correlation matrix of leaf traits, averaged for different plant species at different altitudes in tropical rainforests across an altitudinal gradient in Sri Lanka.

| Pearson Correlation Coefficients (*r*), n = 20<br>Prob > \|*r*\| under H0: *r* = 0 |  |  |  |  |  |  |  |  |  |  |
| --- | --- | --- | --- | --- | --- | --- | --- | --- | --- | --- |
| Trait | GCL | SD | ED | SI | PCI | LBA | LMA | N | P | N:P |
| †GCL | 1.00000 | **-0.792**<br>**<.0001‡** | **-0.542**<br>**0.0135** | -0.214<br>ns | **0.706**<br>**0.0005** | 0.177<br>ns | -0.329<br>ns | -0.371<br>ns | -0.391<br>ns | 0.068<br>ns |
| SD | - | 1.000 | **0.681**<br>**0.0009** | 0.212<br>ns | -0.211<br>ns | -0.132<br>ns | -0.086<br>ns | 0.368<br>ns | 0.258<br>ns | 0.147<br>ns |
| ED | - | - | 1.000 | **-0.504**<br>**0.0234** | -0.114<br>ns | 0.114<br>ns | -0.056<br>ns | **0.477**<br>**0.0333** | 0.358<br>ns | -0.114<br>ns |
| SI | - | - | - | 1.000 | -0.062<br>ns | -0.085<br>ns | 0.087<br>ns | -0.279<br>ns | -0.166<br>ns | 0.190<br>ns |
| PCI | - | - | - | - | 1.000 | 0.352<br>ns | **-0.566**<br>**0.0093** | -0.360<br>ns | -0.291<br>ns | 0.067<br>ns |
| LBA | - | - | - | - | - | 1.00000 | -0.328<br>ns | -0.103<br>ns | 0.226<br>ns | -0.357<br>ns |
| LMA | - | - | - | - | - | - | 1.000 | -0.162<br>ns | -0.065<br>ns | -0.176<br>ns |
| N | - | - | - | - | - | - | - | 1.000 | **0.612**<br>**0.0042** | -0.007<br>ns |
| P | - | - | - | - | - | - | - | - | 1.000 | **-0.582**<br>**0.0071** |
| N:P | - | - | - | - | - | - | - | - | - | 1.000 |

‡Significant correlations are shown in bold.
†GCL - Guard cell length; SD – Stomatal density; ED – Epidermal density; SI – Stomatal index; PCI – Potential conductance index; LBA – Leaf blade area; LMA – Leaf mass per area; N – Leaf nitrogen concentration; P – Leaf phosphorus concentration; N:P – Leaf N:P ratio.

There was a highly-significant (p<0.01) negative correlation between LMA and PCI. Among the leaf nutrients, N and P were positively correlated while leaf N:P was negatively correlated to leaf P. Species means of ‘PNUE’ and ‘PPUE’ were negatively correlated to LMA (Fig. 5.a, b). Strength of these correlations increased when species-level variation within each genus was removed by averaging across species at each altitude (Fig. 5.c, d).

**Figure 5.**
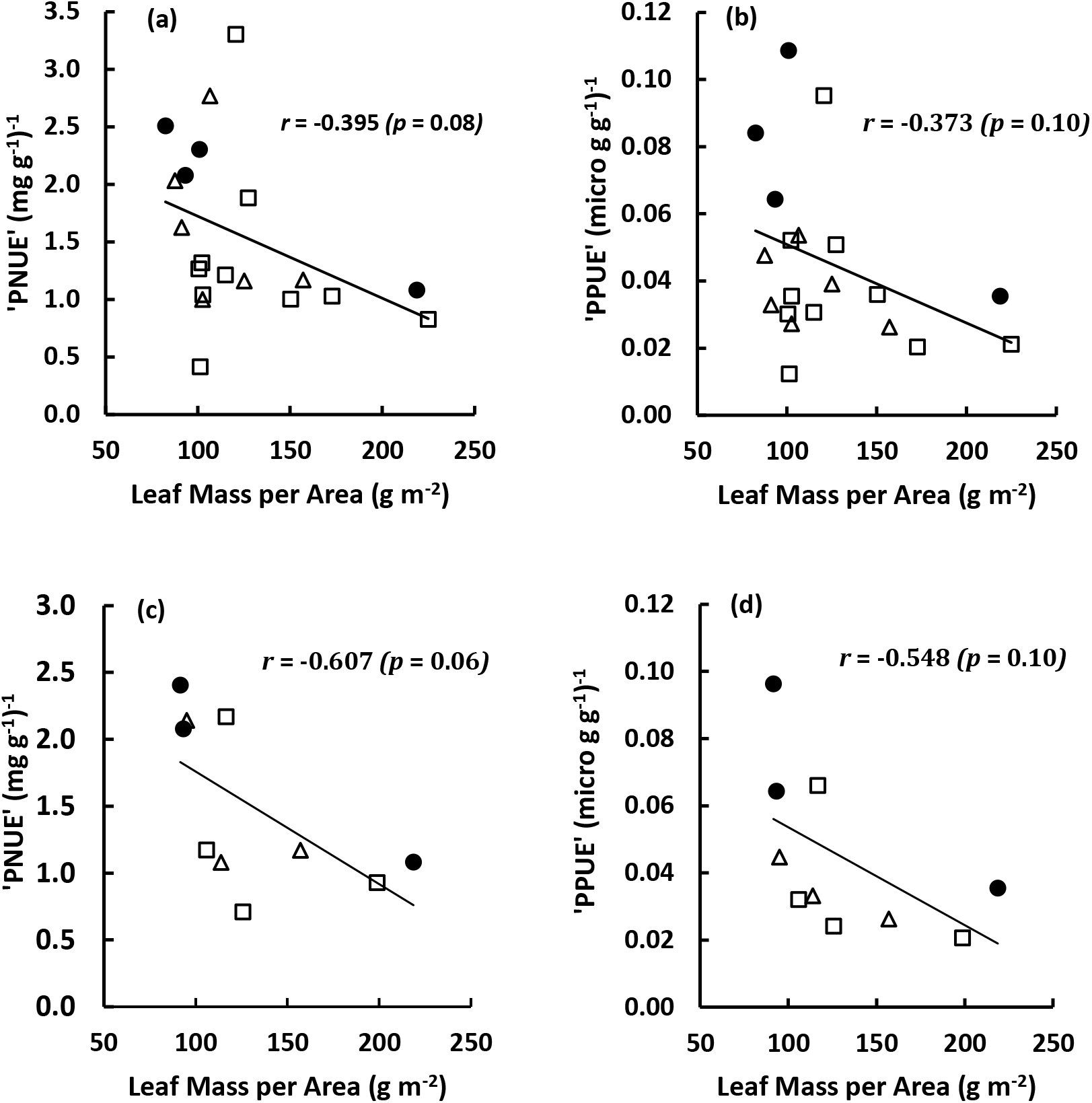
Correlation of proxy indices of photosynthetic nitrogen use efficiency (‘PNUE’)’ and phosphorus use efficiency (‘PPUE’)’ to leaf mass per area (LMA) of different plant species (a, b) and genera (c, d) in tropical rainforests in Sri Lanka across an altitudinal range from 150 m to 2100 m above sea level. ● – *Calophyllum*; Δ – *Semecarpus*; □ – *Syzygium*. ‘PNUE’ and ‘PPUE’ were calculated as the respective ratios of potential conductance index (unitless) and leaf N and P concentrations.

### Trends with key climatic variables

When the data from all species and altitudes were pooled, stomatal density (SD), guard cell length (GCL), epidermal density (ED) and stomatal index (SI) did not show significant (p>0.05) linear relationships with long-term mean air temperature (T_AV_), annual precipitation (R_F_) and daily solar irradiance (S_R_). However, potential conductance index (PCI) showed highly-significant (*p*<0.001) positive linear relationships with T_AV_, R_F_ and S_R_ (Fig. 6.a-c). Leaf mass per area (LMA) showed significant (*p*<0.01) negative linear relationships with T_AV_, R_F_ and S_R_ (Fig. 6.d-f).

**Figure 6.**
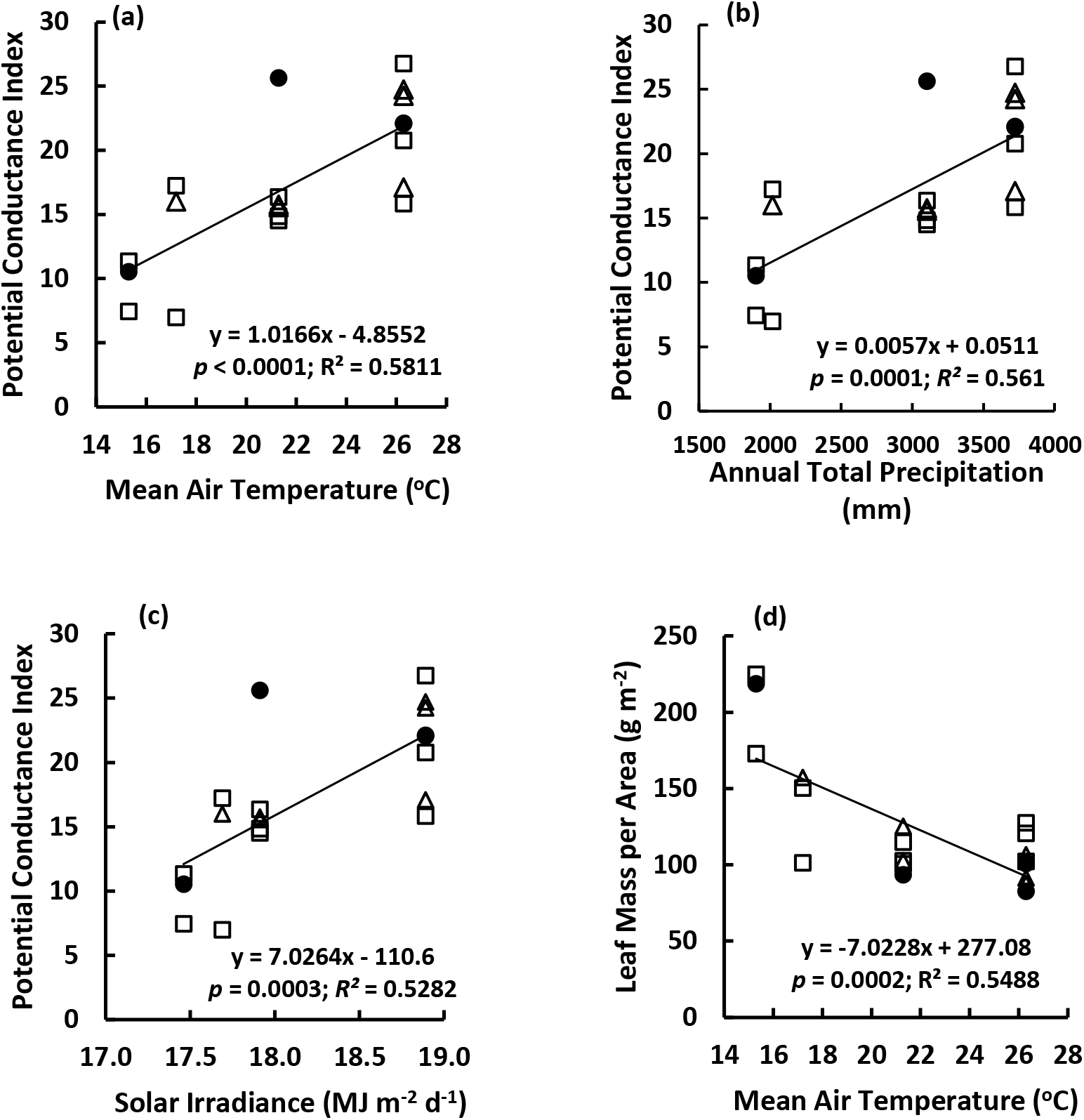
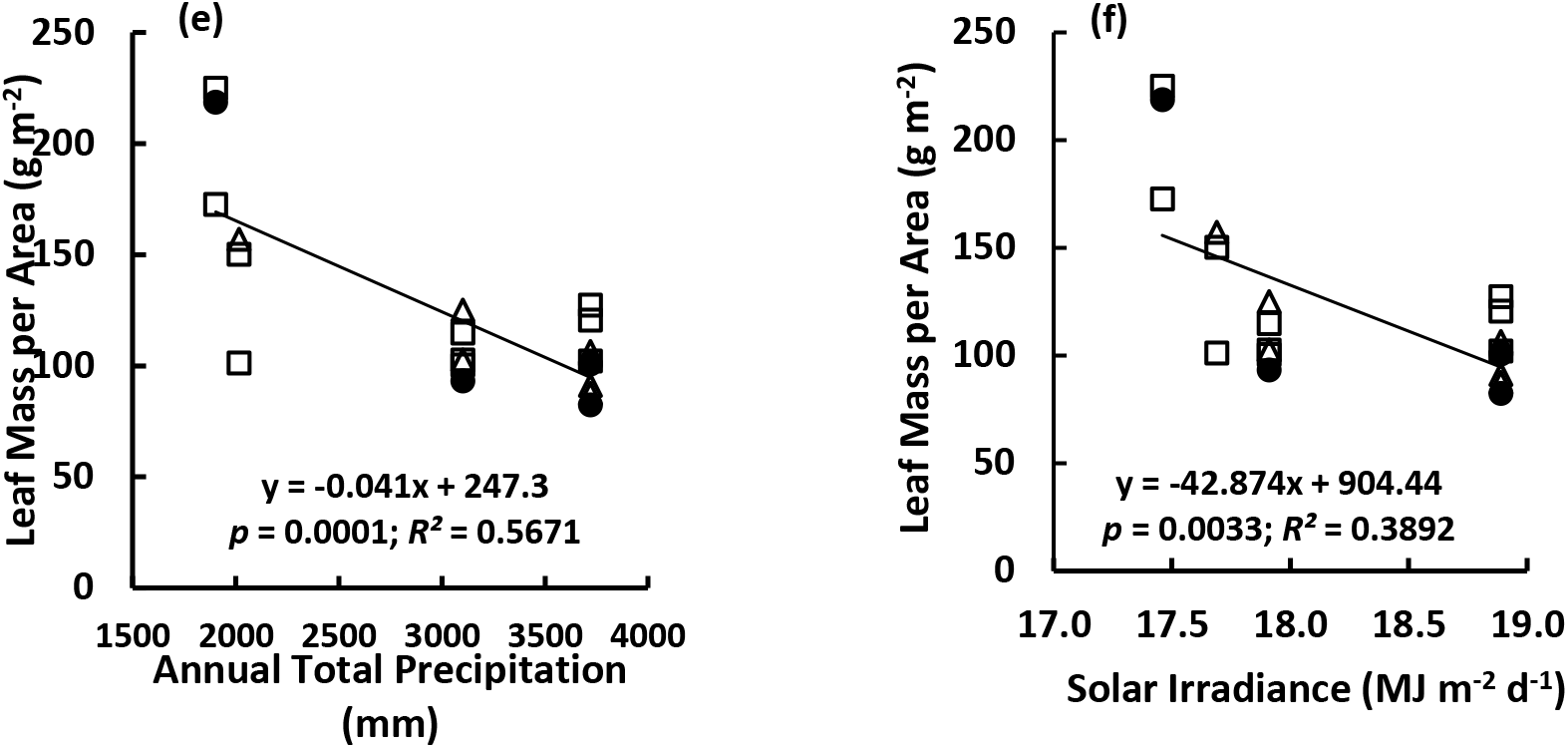
Variation of potential conductance index (a-c) and leaf mass per area (d-f) of different plant species with temperature (a, d), precipitation (b, e) and solar irradiance (c, f) in tropical rainforests in Sri Lanka across an altitudinal range from 150 m to 2100 m above sea level. ● – *Calophyllum*; Δ – *Semecarpus*; □ – *Syzygium*. Each data point is a species-level mean value.

Because of the substantial variance component due to the random effect of genus (Table 5), relationships of LBA with climatic variables were examined separately for *Semecarpus* which had substantially higher LBA than *Calophyllum* and *Syzygium* (Fig. 3.b), whose LBA trends were examined together. Mean LBA of different *Semecarpus* species did not show significant trends with any of the climate variables (Fig. S5). However, when LBA of *Semecarpus* species at each altitude were averaged, the genus-level mean LBA of *Semecarpus* showed clear increasing trends with T_AV_, R_F_ and S_R_ (Fig. S6). Pooled species-level and genus-level LBA means of *Syzygium* and *Calophyllum* showed highly-significant (*p*<0.01) increasing trends with T_AV_, R_F_ and S_R_.

Leaf N and P showed second-order polynomial trends with T_AV_, R_F_ and S_R_ (Fig. 7). Trends for leaf P showed greater scatter (i.e. lower *R^2^* values and greater *p*) than those for leaf N. Leaf N:P ratio did not show significant trends with any climate variable (data not shown). Proxy indices of photosynthetic N and P use efficiencies (‘PNUE’ and ‘PPUE’) showed highly-significant positive trends with T_AV_, R_F_ and S_R_ (Fig. 8).

**Figure 7.**
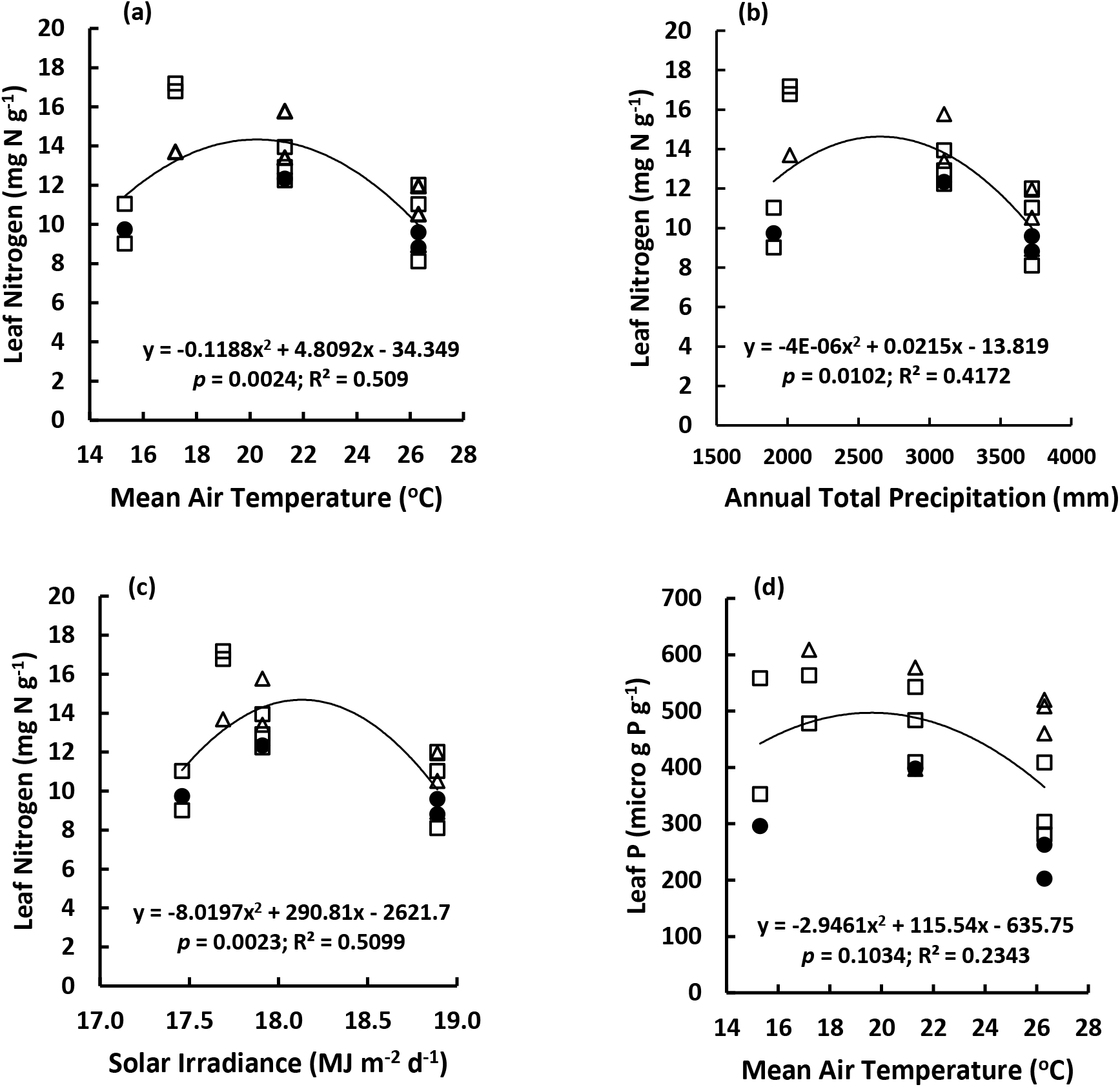
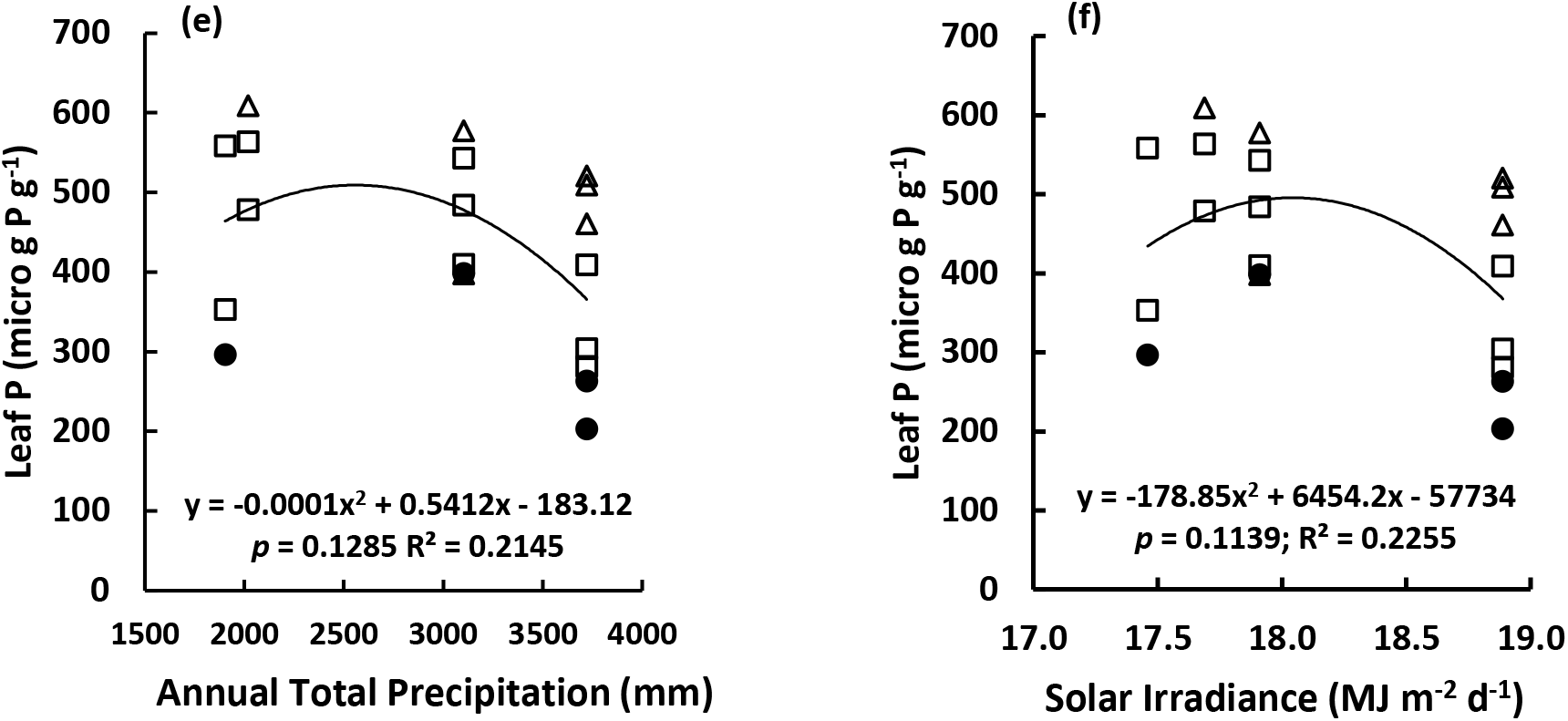
Variation of leaf nitrogen (a – c) and phosphorus (d – f) concentrations of different plant species with temperature (a, d), precipitation (b, e) and solar irradiance (c, f) in tropical rainforests in Sri Lanka across an altitudinal range from 150 m to 2100 m above sea level. ● – *Calophyllum*; Δ – *Semecarpus*; □ – *Syzygium*. Each data point is a species-level mean value.

**Figure 8.**
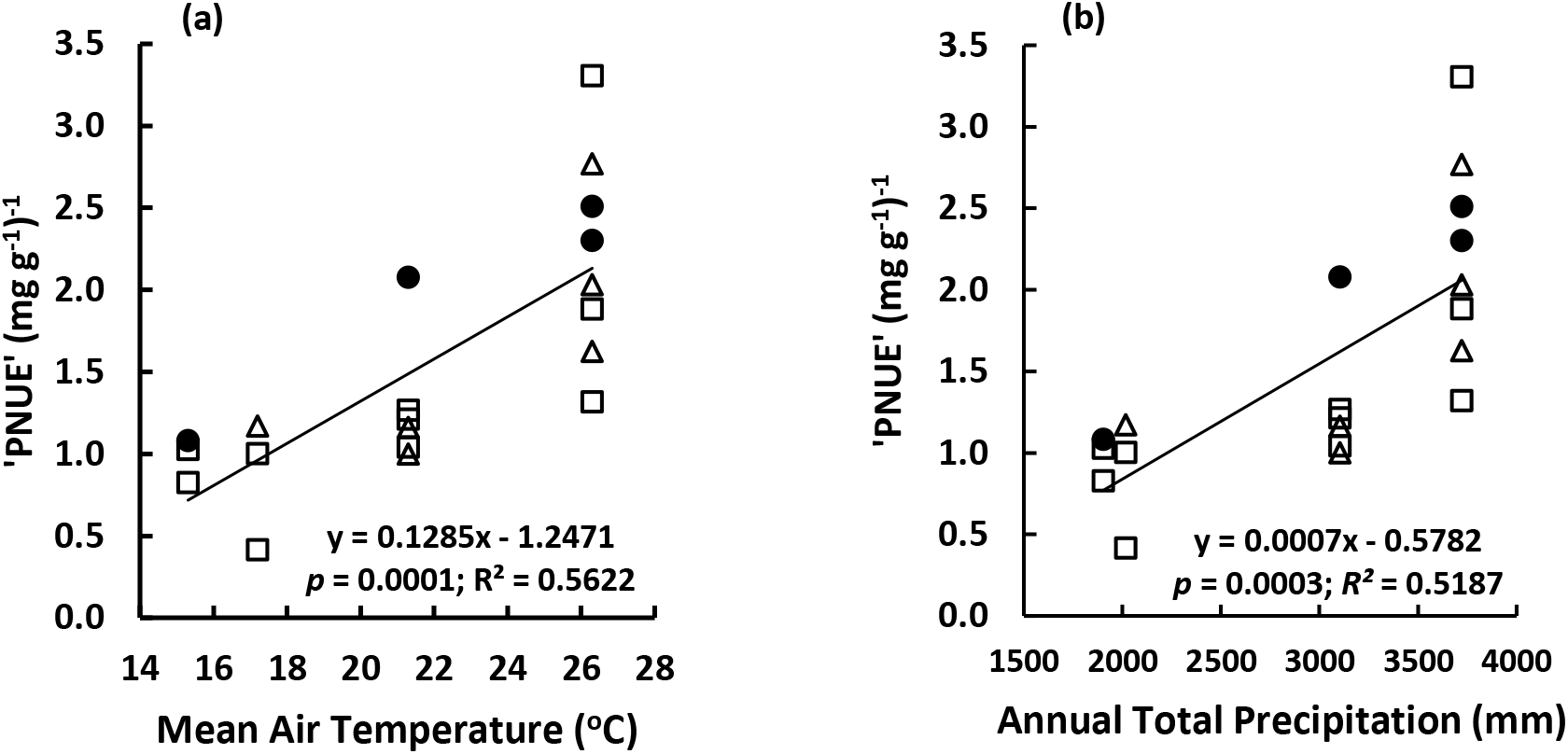
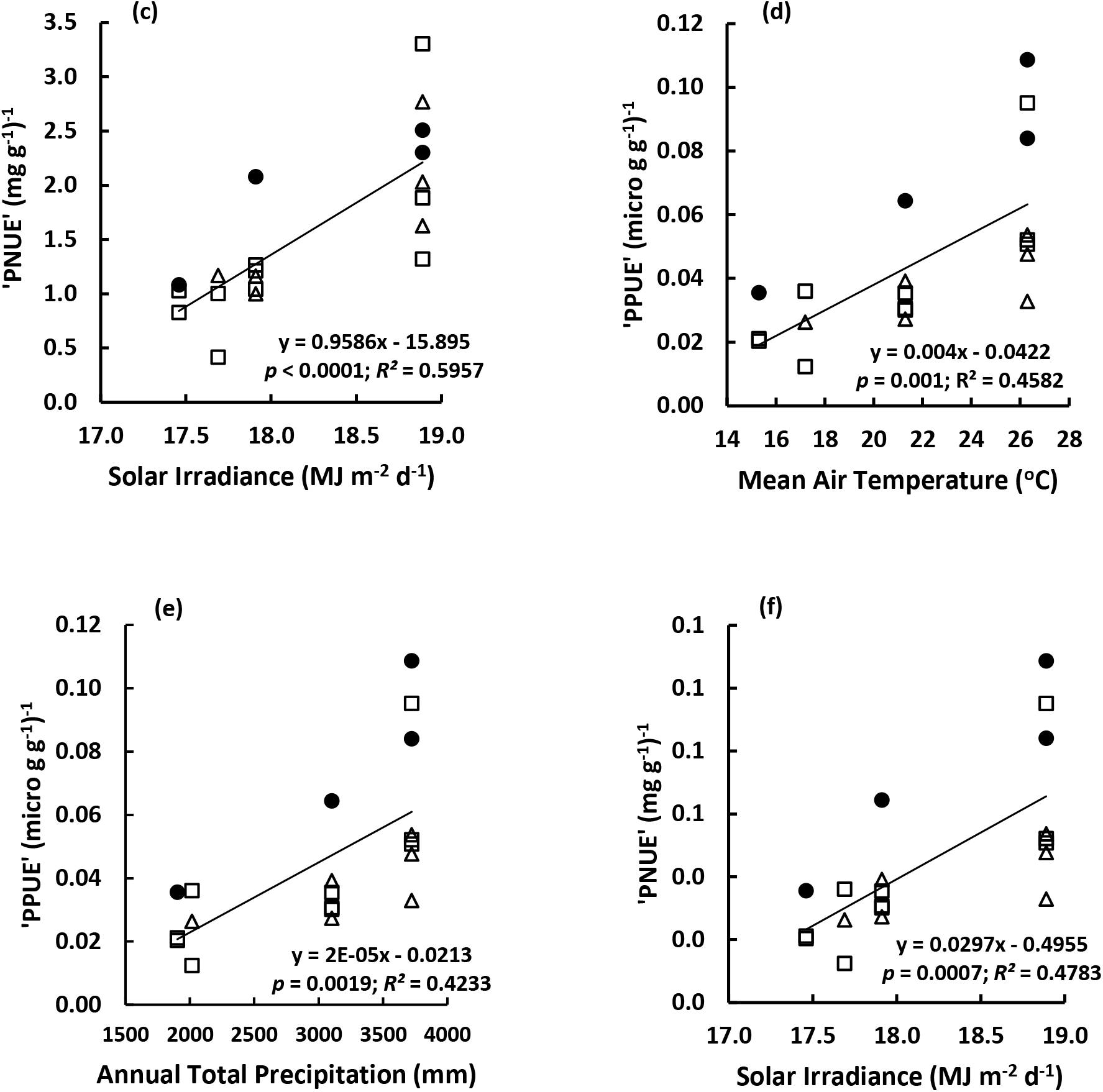
Variation of proxy indices of photosynthetic nitrogen use efficiency (‘PNUE’)’ and phosphorus use efficiency (‘PPUE’)’ of different plant species with temperature (a, d), precipitation (b, e) and solar irradiance (c, f) in tropical rainforests in Sri Lanka across an altitudinal range from 150 m to 2100 m above sea level. ● – *Calophyllum*; Δ – *Semecarpus*; □ – *Syzygium*. Each data point is a species-level mean value.

### Factor analysis

Four factors that had Eigen values greater than one were extracted initially and these accounted for 82% of the total variance in leaf traits in different plant species. The rotated factor pattern, following a varimax rotation, and the variables which loaded to each factor are given in Table 7.

**Table 7.** Rotated factor pattern for extracted factors in a factor analysis of leaf traits of plant species distributed along an altitudinal gradient in tropical rainforests in Sri Lanka.

| Variable | Loadings<sup>†</sup> of variables to factors |  |  |  |
| --- | --- | --- | --- | --- |
|  | Factor1 | Factor2 | Factor3 | Factor4 |
| Stomatal density (SD) | 94* | 3 | 11 | 16 |
| Epidermal density (ED) | 73 | 9 | -11 | -48 |
| Leaf nitrogen (N) | 60 | -9 | -10 | -52 |
| Guard cell length (GCL) | -83* | 47 | 11 | -18 |
| Potential conductance index (PCI) | -34 | 84* | 7 | 2 |
| Leaf mass per area (LMA) | -14 | -86* | -6 | 10 |
| Leaf N:P ratio | 10 | 14 | 90* | 9 |
| Leaf blade area (LBA) | -6 | 56 | -63 | 9 |
| Leaf phosphorus (P) | 45 | -9 | -72 | -24 |
| Stomatal index (SI) | 9 | -7 | 10 | 94* |

<sup>†</sup>Multiplied by 100; \*Loadings greater than 0.8.

A variable was considered to load on a factor if its loading on that factor is greater than 0.8 and if the same variable does not load on another factor. Accordingly, two variables, namely, SD and GCL, loaded on Factor 1, which accounted for 33% of the total variance of leaf traits. Leaf mass per area and PCI loaded on Factor 2 (21% of total variance) while leaf N:P (16%) and SI (12%) loaded on Factors 3 and 4 respectively.

Figure 9 shows the distribution of different tree species in the factor space based on their rotated factor scores for Factors 1 and 2. It can be observed that the three species found at 2100 m in Pidurutalagala are located away from the rest (group A). Two of the three species at 1800 m in Hakgala are located together (group B), but one (*Semecarpus obovata*) is located away from all species in all locations. Out of the six species found at 1050 m in Sinharaja-Enasalwatte, the three *Syzygium* species are located close to each other (group C). However, the two *Semecarpus* species and the *Calophyllum* species are located away from the cluster formed by the three *Syzygium* species and from each other. A majority of the eight species found at 150 m in Kanneliya have formed two separate groups. One group (D) is formed by the two *Calophyllum* species (i.e. *C. bracteatum* and *C. cordata-oblongum*). Notably, *Calophyllum acidus* from Sinharaja-Enasalwatte also is located within this species group. All three *Semecarpus* species in Kanneliya are located within group E. Interestingly, *Semecarpus gardneri* in Sinharaja-Enasalwatte also is located in group E so that *S. gardneri* in both locations is within the same species group. Two of the three *Syzygium* species in Kanneliya (*S. alubo* and *S. firmum*) are also located within group E along with the three *Semecarpus* species. Notably, the other *Syzygium* species in Kanneliya (*S. neesianum*) is located close to group C, which consists of the three *Syzygium* species in Sinharaja-Enasalwatte. *Semecarpus obovata* (at Hakgala) and *S. parvifolia* (at Sinharaja-Enasalwatte) which are located away from all other species at all locations form a final species group (F).

**Figure 9.**
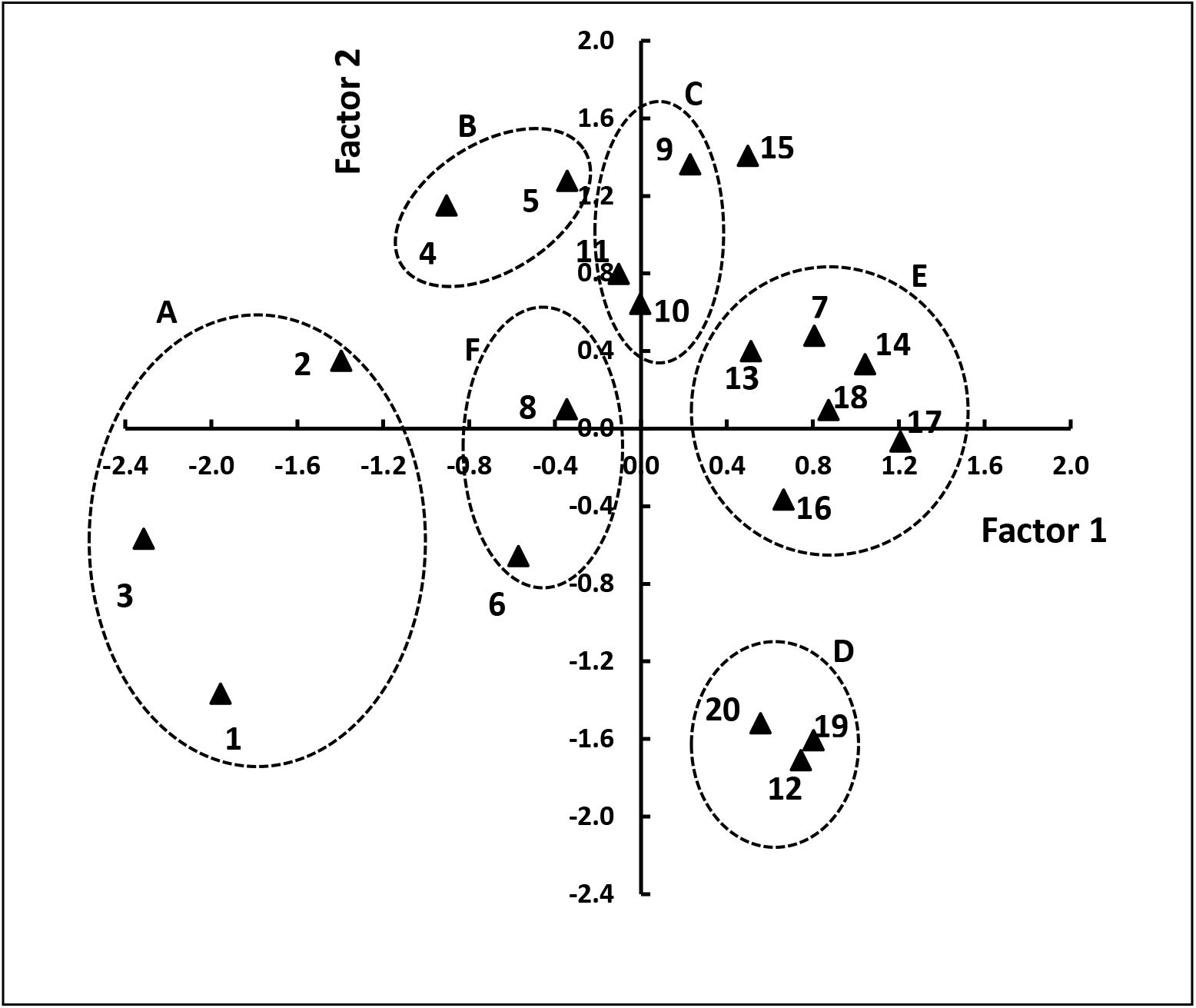
Distribution of plant species in factor space based on factor scores (after rotation) of Factors 1 and 2. 1 – 3: Species at 2100 m in Pidurutalagala; 4 – 6: Species at 1800 m in Hakgala; 7 – 12: Species at 1050 m Sinharaja-Enasalwatte; 13 -20: Species at 150 m in Kanneliya. 1: *Calophyllum walkeri*; 2: *Syzygium rotundifolium*; 3: *Syzygium revolutum*; 4: *Syzygium zeylanicum*; 5: *Syzygium paniculatum*; 6: *Semecarpus obovata*; 7: *Semecarpus gardneri* (At Sin-Enasalwatte); 8: *Semecarpus parvifolia*; 9: *Syzygium cylindricum*; 10: *Syzygium micranthum*; 11: *Syzygium spp.*; 12: *Calophyllum acidus*; 13: *Syzygium alubo*; 14: *Syzygium firmum*; 15: *Syzygium neesianum*; 16: *Semecarpus gardneri* (At Kanneliya); 17: *Semecarpus subpeltata*; 18: *Semecarpus walkeri*; 19: *Calophyllum bracteatum*; 20: *Calophyllum cordato-oblongum*.

### Cluster analysis

The dendrogram produced by cluster analysis of the leaf traits of different plant species using the complete linkage method is shown in Fig. 10. At a linkage distance of just above 0.5, six species clusters were identified. Multivariate analysis of variance (MANOVA) showed that the six clusters were significantly different (Wilks’ Lambda Statistic = 0.00000956, p<0.0001). Notably, all four *Calophyllum* species across the altitudinal range from 150 m to 2100 m grouped in to Cluster 2. Cluster 1 consists of two sub-clusters, one having two *Semecarpus* species and the other having three *Syzygium* species. It is notable that the species in both these sub-clusters come from different altitudes. Cluster 3 has two *Syzygium* species from 2100 m and 1800 m, along with a *Semecarpus* species from 150 m. Cluster 4 consists of two *Syzygium* species, one from 1050 m and the other from 150 m. Cluster 5 has three *Syzygium* species from three different altitudes (i.e. 1800 m, 1050 m and 150 m) and one *Semecarpus* species from 1050 m. Lastly, Cluster 6 has two *Semecarpus* species from 150 m. Therefore, while all *Calophyllum* species have grouped into one cluster, there has been mixing between *Syzygium* and *Semecarpus* species during clustering. In all three genera, species from different altitudes have come together in common clusters.

**Figure 10.**
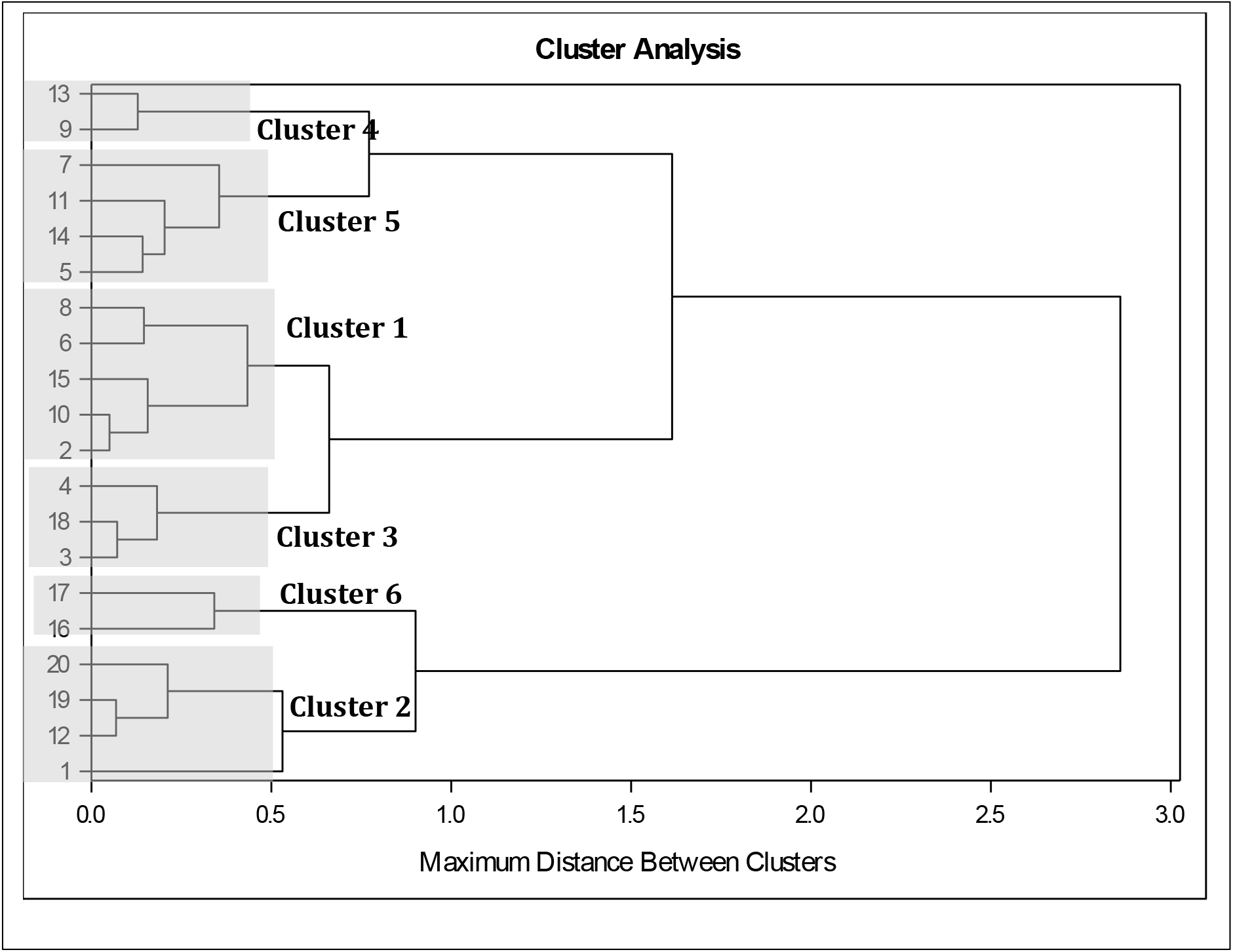
Dendrogram of cluster analysis based on leaf traits of plant species from three genera in permanent sampling plots within tropical rainforests along an altitudinal gradient in Sri Lanka. 1 – 3: Species at 2100 m asl in Pidurutalagala; 4 – 6: Species at 1800 m in Hakgala; 7 – 12: Species at 1050 m in Sinharaja-Enasalwatte; 13 -20: Species at 150 m in Kanneliya. 1: *Calophyllum walkeri*; 2: *Syzygium rotundifolium*; 3: *Syzygium revolutum*; 4: *Syzygium zeylanicum*; 5: *Syzygium paniculatum*; 6: *Semecarpus obovata*; 7: *Semecarpus gardneri* (In Sin-Enasalwatte); 8: *Semecarpus parvifolia*; 9: *Syzygium cylindricum*; 10: *Syzygium micranthum*; 11: *Syzygium* sp.; 12: *Calophyllum acidus*; 13: *Syzygium alubo*; 14: *Syzygium firmum*; 15: *Syzygium neesianum*; 16: *Semecarpus gardneri* (In Kanneliya); 17: *Semecarpus subpeltata*; 18: *Semecarpus walkeri*; 19: *Calophyllum bracteatum*; 20: *Calophyllum cordato-oblongum*. Six clusters are identified by hatched boxes.

## Discussion

We discuss the above results in terms of how they have provided answers to the questions that we posed at the beginning of this work.

### Do the altitudinal responses of stomatal traits vary for different taxa?

As the genus x altitude interaction did not account for a substantial variance component in any of the stomatal traits (Table 4), our results do not support the hypothesis that the altitudinal variation of individual stomatal anatomical traits is different among different taxa in TRFSL. Out of the stomatal traits, only PCI, a composite trait combining the effects of stomatal size (GCL) and density (SD) in determining leaf gas exchange capacity, and a proxy for photosynthetic capacity (Wong et al. 1979, Farquhar and Sharkey 1982) showed variation with altitude (Table 3). It is notable that neither the genus nor genus x altitude interaction contributed appreciably to the total variance of PCI, thus indicating that the observed altitudinal response of PCI is common for different taxa. Interestingly, the two component traits of PCI (i.e. GCL and SD) show substantial variation among genera (Table 4) while not responding to altitudinal variation. Therefore, our observation of a common altitudinal response of PCI supports a hypothesis that the genetically diverse altitudinal responses of individual stomatal traits are part of an integrated altitudinal response in the leaf gas exchange capacity.

### Are the altitudinal responses of different leaf traits linked in an integrated/coordinated response?

Among the other leaf traits, LMA, LBA and leaf N showed significant altitudinal trends (Table 3). Despite LBA showing an appreciable variance component due to genus, the altitudinal trends of the above three traits did not show variation among the three genera (Table 4, Figs. 3, 4). This affirmed the absence of evidence to support Hypothesis 1 with regard to leaf structural traits and key nutrients. However, two of our observations support the hypothesis that the altitudinal responses of different leaf traits are interlinked across different taxa to form a coordinated response across different taxa) (Hypothesis 2). One is the significant negative correlation between PCI and LMA (Table 6). The other is the result from factor analysis where PCI and LMA formed one of the two major variable constructs that explained the observed variance of leaf traits (Table 7). The negative correlation between PCI and LMA is mechanistically plausible. Increasing LMA indicates greater investment of biomass per unit leaf area, especially in structural tissue, thus enabling a longer leaf lifespan. The leaf economic spectrum (Wright et al. 2004, Onoda and Wright 2018) specifies that greater investment in leaf structure is accompanied by lower investment in photosynthetic machinery, which entails a ‘slower return’ in terms of assimilate production. Lower photosynthetic capacity does not require a high CO_2_ absorption capacity via a higher stomatal conductance. As PCI can be considered as the maximum possible stomatal conductance (g_max_), it is physiologically plausible that photosynthetic capacity and g_max_ are negatively correlated to greater investment in leaf structural tissue (i.e. higher LMA) and longer leaf lifespan.

We find only partial evidence that variation of N and P across altitudes, their associated climatic parameters and genera is part of a coordinated response along with stomatal anatomical and leaf structural traits. For example, we could expect leaf N to be positively correlated to PCI if higher leaf N has resulted in greater investment of N in photosynthetic machinery (e.g. Rubisco). Similarly, a positive correlation could be expected between PCI and leaf P, if higher leaf P indicates greater investment in energy storage molecules such as ATP and NADPH. Across the whole range of altitudes in our study, leaf N and P were not significantly correlated to PCI (Table 7). However, both leaf N and P, which had shown increases from low (150 m) to mid (1050 m) altitude, showed decreases from mid- to high (> 1800 m) altitudes (Fig. 4) where parallel decreases have occurred in PCI as well (Fig. 2). Therefore, the reductions in N and P from mid- to high altitudes could be due to down-regulation of photosynthetic capacity, as shown by decreased PCI. Based on global scale studies, leaf N and P were also expected to be negatively correlated to LMA as greater investment in leaf structural tissue in the high-LMA leaves would mean reduced allocation of N and P to leaves. However, across the whole range of altitudes we have not found significant correlations between LMA and leaf nutrients. On the other hand, the expected strong negative correlations are found between LMA and leaf nutrients from mid- to high altitudes. Therefore, our results suggest that leaf nutrients vary as part of integrated/coordinated leaf trait response at lower temperature and rainfall regimes (i.e. mid- to high altitudes), but not at higher temperature and rainfall regimes (i.e. low altitudes). Shifts in trait relationships with variation in climate have been observed by (Wright et al. 2005).

Examination of trait relationships and climate responses of nutrient use efficiencies of our data provide evidence for leaf nutrients also being part of a coordinated response in concert with stomatal anatomical and leaf structural traits. Both ‘PNUE’ and ‘PPUE’ showed negative correlations with LMA (Fig. 8). As suggested by Onoda and Wright (2018), allocation of a higher proportion of leaf N and P to structural tissue could have caused these reductions in ‘nutrient use efficiencies’ with LMA. Furthermore, the positive relationships of ‘PNUE’ and ‘PPUE’ with T_AV_, R_F_ and S_R_ (Fig. 8) are in accordance with a strategy of increased investment of N and P in photosynthetic machinery in higher temperature and precipitation environments where LMA is lower and photosynthetic capacity is higher (Fig. 6).

### How strong is the influence of climate variation across altitudinal gradients on responses of leaf traits to altitude?

Assessing the influence of climatic variation across the altitudinal gradient on leaf traits and their associations is constrained in our work by the limited number of data points (i.e. four altitudes). However, the strong trends of PCI (positive, Figs. 6.a-c) and LMA (negative, Figs. 6.d-f) with temperature (T_AV_), precipitation (R_F_) and incident solar radiation (S_R_) indicate significant environmental control over key leaf traits. The absence of a genus level effect as shown by its very low variance component (Tables 4, 5) show that climatic factors exert an influence that overrides genetic variation. Mechanistic explanations are available for the observed trends between the above leaf traits and climatic factors. Increase of PCI with increasing R_F_ agrees with the tight control exerted by water availability on leaf gas exchange capacity. Species in sites of higher precipitation have evolved greater gas exchange capacity (i.e. higher PCI), probably to maximize carbon assimilation without being constrained by the possibility of water stress due to excessive transpiration. It is possible that the higher PCI at higher R_F_ sites is the result of a combination of climatic factors that favour higher assimilation rates. This is because in the present study, higher R_F_ at lower altitudes (e.g. 150 m) is accompanied by higher T_AV_, S_R_ and pCO_2_, as can be assumed with higher WVP, (Table 1), all of which promote photosynthesis. In contrast, as R_F_ declined progressively with increasing altitude, the lower R_F_ sites are at higher altitudes where T_AV_, S_R_ and pCO_2_ also are lower. The lower T_AV_, T_min_ and T_max_ levels at higher altitudes of the present study are more likely than their R_F_ to impose limitations on photosynthesis (James et al. 1994, Cabrera et al. 1998, Zhang et al. 2005). Therefore, it is possible that the lower PCI at the lower R_F_ sites have occurred more as a response to decreased photosynthetic potential at lower temperatures than a response to lower R_F_.

The negative trends of LMA with T_AV_, R_F_ and S_R_ (Fig. 6) indicate increased allocation of leaf biomass per unit leaf area. While the negative trend with R_F_ agrees with the broader global trend, the decrease of LMA with increasing T_AV_ is opposite to the trend shown with a broader range of plant species and climates (Wright et al. 2004, 2005). However, when the trend for evergreens is separated from the global trend (Wright et al. 2005), it agrees with that observed in our study. It is possible that the increase of LMA with decreasing R_F_ and T_AV_ is a consequence of restrictions imposed by both these climatic variables on LBA. In all three genera, LBA decreased with decreasing R_F_ and T_AV_ (Figs. S5, S6) probably because of their effects on leaf cell division (CDR) and expansion rates (CER). There is ample evidence from agricultural crops that in plant tissues growing without water deficits, both leaf CDR and CER increases with increasing temperature above a base temperature (Squire and Ong 1983, Squire 1990, Chapman et al. 1993, Salah and Tardieu 1996, Tardieu et al. 2005). Furthermore, CDR and CER are highly-sensitive to water availability, showing reduction with water deficit (Squire et al. 1983, Ong et al. 1985, Granier and Tardieu 1999, Tardieu et al. 2000, 2005, Chenu et al. 2008). It is highly-likely that these effects on CDR and CER have combined to decrease LBA with decreasing T_AV_ and R_F_ along the altitudinal gradient in our study (Figs. S5, S6). Such a restriction of LBA would concentrate the biomass allocated to leaves within a smaller area, thus causing the observed increase of LMA with decreasing T_AV_ and R_F_. This increase of LMA is in agreement with the observed trend of thicker and leathery leaves in environments with lower R_F_ and with longer leaf lifespans (LL) in cooler and drier environments (Niinemets 2001, Wright et al. 2001, 2004). This trend is also in accordance with an integrated response, which includes the observed reduction of PCI and photosynthetic capacity with decreasing T_AV_, R_F_ and S_R_. Even though photosynthesis was not measured in the present study, mass-based photosynthetic capacity has been shown to decline with increasing LMA (Osnas et al. 2013, Onoda and Wright 2018). Therefore, the highly-significant negative correlation between PCI and LMA observed in our study (Table 6) is mechanistically plausible and is in agreement with the global trends.

The highly-significant positive correlation between leaf N and P is in agreement with global scale observations (Wright et al. 2004, Reich and Oleksyn 2004, Osnas et al. 2013, Onoda and Wright 2018). The second-order polynomial trends of leaf N and P with T_AV_ (Fig. 7) are similar to the global-scale patterns for a wider range of species and climates (Reich and Oleksyn 2004). The absence of a response to climatic factors in leaf N:P ratio indicates that the responses of N and P to climate variation occurs in a way that N:P is conserved across different climates. This is confirmed by N:P being extracted as one of the underlying variable constructs influencing the variation of leaf traits across species, genera and climates.

### Evidence from factor analysis for an integrated/coordinated altitudinal response

We preferred to perform a factor analysis instead of a principal component analysis because of our assumption based on previous work (Wright et al. 2004, Onoda and Wright 2018) that there is an underlying structure of trait assemblies (i.e. underlying variable constructs) that causes the observed variation of leaf traits across altitudes, genera and species (Child 1990, Suhr 2005). The four extracted factors, which accounted for 82% of the variance of measured leaf traits, represented physiologically-meaningful trait assemblies. Factor 1, composed of SD and GCL, constituted an underlying composite factor incorporating two properties that determine the abundance of stomata on the leaf surface and their size. Factor 2, being loaded by PCI and LMA, clearly constituted the leaf economic spectrum (Wright et al. 2004). Factor 3 constituted an underlying factor based on leaf nutrient contents, the leaf N:P ratio. Stomatal index, a measure of the fraction of stomata relative to the total cell population on the leaf surface, formed the fourth underlying factor. The distribution of different plant species and genera in the factor space of Factors 1 and 2 did not show a clear separation of species or genera. This confirms that the influence of underlying factors runs across taxonomic boundaries. This is confirmed by the clustering of species in cluster analysis, where except for the four *Calophyllum* species, there was considerable mixing of species of *Syzygium* and *Semecarpus* from different altitudes.

## Conclusions

We, therefore, conclude that: (a) the response of stomatal anatomy to altitude does not differ among taxa of trees in TRFs; (b) the altitudinal responses of leaf structural traits and major nutrient concentrations are interlinked in a coordinated response across different taxa; and (c) climate exerts a strong influence on the variation of leaf traits to environmental gradients, which spans across taxonomic boundaries of species and genera. We also find evidence that the coordinated response of leaf traits to climate variation is in accordance with the leaf economics spectrum. These findings have important implications on how leaf traits and their responses to climate variations are represented in dynamic vegetation models to predict the response of TRFs to future climate change.

## Supplementary data

Supplementary data of this article is available separately.

## Acknowledgements

We thank the Departments of Forest Conservation and Wildlife Conservation, Sri Lanka for logistical support, Gemunu Wijesooriya for assistance in leaf nutrient analysis and Chameesha Madhumali, Dineth Dhanushka and Suneth Kanishka for field assistance.

## Authors’ contributions

JC, IC and BW conceptualized and designed the work. NS and DS established permanent sampling plots and carried out the plant census, with assistance from HJ and AW in taxonomic identification and description. BW collected leaf samples and carried out all measurements under the supervision of IC. JC, IC and BW did the statistical analysis of data. JC wrote the paper with assistance from IC, AW and BW. IC and BW reviewed the initial draft and IC reviewed the final draft.

## List of supplementary figures

**Figure S1.** Box plots of distributions of stomatal traits of plant species of tropical rainforests in Sri Lanka at different altitudes: (a) guard cell length; (b) stomatal density; (c) epidermal density; (d) stomatal index; (e) potential conductance index. Abbreviations for species: Calbrac – *Calophyllum bracteatum*; Calcoobl – *Calophyllum cordato-oblongum*; Semgard – *Semecarpus gardneri* (at 150 and 1050 m); Semsub – *Semecarpus subpeltata;* Semwalk – *Semecarpus walkeri*; Syzalu – *Syzygium alubo*; Syzferm – *Syzygium firmum*; Syznees – *Syzygium neesianum*; Calcid – *Calophyllum acidus*; Semparv – *Semecarpus parvifolia*; Syzcyl – *Syzygium cylindricum*; Syzmic – *Syzygium micranthum*; Syznew – *Syzygium* sp.; Semobo – *Semecarpus obovata*; Syzpan – *Syzygium paniculatum*; Syzzel – *Syzygium zeylanicum*; Calwalk – *Calophyllum walkeri*; Syzrev – *Syzygium revolutum*; Syzrot – *Syzygium rotundifolium*.

**Figure S2.** Box plots of distributions of leaf structural traits of plant species of tropical rainforests in Sri Lanka at different altitudes: (a) leaf mass per area; (b) leaf blade area. Abbreviations for species are as in Fig. S.1.

**Figure S3.** Box plots of distributions of leaf nutrients of plant species of tropical rainforests in Sri Lanka at different altitudes: (a) leaf nitrogen; (b) leaf phosphorus; (c) leaf nitrogen:phosphorus ratio. Abbreviations for species are as in Fig. S.1.

**Figure S4.** Box plots of distributions of leaf nutrients of plant genera of tropical rainforests in Sri Lanka at different altitudes: (a) leaf nitrogen; (b) leaf phosphorus; (c) leaf nitrogen:phosphorus ratio.

**Figure S5.** Species-level variation of leaf blade area (LBA) with climatic variables: (a) temperature; (b) precipitation; (c) solar irradiance. Regression lines are fitted for *Semecarpus* species separately and for *Syzygium* and *Calophyllum* species together. Each data point is a species mean.

**Figure S6.** Genus-level variation of leaf blade area (LBA) with climatic variables: (a) temperature; (b) precipitation; (c) solar irradiance. Regression lines are fitted for *Semecarpus* separately and for *Syzygium* and *Calophyllum* together. Each data point is the mean of different species of a given genus.

## References

Aiba S-I, Kitayama K (1999) Structure, composition and species diversity in an altitude-substrate matrix of rain forest tree communities on Mount Kinabalu, Borneo. Plant Ecol 140:139–157.

Angiosperm Phylogeny Group (2009) An update of the Angiosperm Phylogeny Group classification for the orders and families of flowering plants: APG III. Bot J Linn Soc 161:105–121.

Ashton PS (1981) Myrtaceae. In: Dassanayake MD, Fosberg FR (eds) A Revised Handbook to the Flora of Ceylon: Volume II. Amerind, New Delhi

Bai Y-J, Chen L-Q, Ranhotra PS, Wang Q, Wang Y-F, Li C-S (2015) Reconstructing atmospheric CO_2_ during the Plio-Pleistocene transition by fossil Typha. Glob Chang Biol 21:874–881.

Barry RG (1992) Mountain Weather and Climate. Psychology Press.

Cabrera HM, Rada F, Cavieres L (1998) Effects of temperature on photosynthesis of two morphologically contrasting plant species along an altitudinal gradient in the tropical high Andes. Oecologia 114:145–152.

Caldera HIU, De Costa WAJM, Woodward FI, Lake JA, Ranwala SMW (2017) Effects of elevated carbon dioxide on stomatal characteristics and carbon isotope ratio of Arabidopsis thaliana ecotypes originating from an altitudinal gradient. Physiol Plant 159:74–92.

Chapman SC, Hammer GL, Palta JA (1993) Predicting leaf area development of sunflower. Field Crops Res 34:101–112.

Chenu K, Chapman SC, Hammer GL, Mclean G, Salah HBH, Tardieu F (2008) Short-term responses of leaf growth rate to water deficit scale up to whole-plant and crop levels: an integrated modelling approach in maize. Plant, Cell & Environment 31:378–391.

Child D (1990) The essentials of factor analysis. Cassell Educational.

Dieleman WIJ, Venter M, Ramachandra A, Krockenberger AK, Bird MI (2013) Soil carbon stocks vary predictably with altitude in tropical forests: Implications for soil carbon storage. Geoderma 204–205:59–67.

Doheny-Adams T, Hunt L, Franks PJ, Beerling DJ, Gray JE (2012) Genetic manipulation of stomatal density influences stomatal size, plant growth and tolerance to restricted water supply across a growth carbon dioxide gradient. Philos Trans R Soc Lond B Biol Sci 367:547–555.

Farquhar GD, Sharkey TD (1982) Stomatal conductance and photosynthesis. Annu Rev Plant Physiol 33:317–345.

Fick SE, Hijmans RJ (2017) WorldClim 2: new 1-km spatial resolution climate surfaces for global land areas. Int J Climatol 37:4302–4315.

Franks PJ, Beerling DJ (2009) Maximum leaf conductance driven by CO_2_ effects on stomatal size and density over geologic time. Proc Natl Acad Sci USA 106:10343–10347.

Givnish TJ (1999) On the causes of gradients in tropical tree diversity. J Ecol 87:193–210.

Granier C, Tardieu F (1999) Water deficit and spatial pattern of leaf development. Variability In responses can Be simulated using a simple model of leaf development. Plant Physiol 119:609–620.

Haworth M, Killi D, Materassi A, Raschi A (2015) Coordination of stomatal physiological behavior and morphology with carbon dioxide determines stomatal control. Am J Bot 102:677–688.

Hofhansl F, Chacón-Madrigal E, Fuchslueger L, Jenking D, Morera-Beita A, Plutzar C, Silla F, Andersen KM, Buchs DM, Dullinger S, Fiedler K, Franklin O, Hietz P, Huber W, Quesada CA, Rammig A, Schrodt F, Vincent AG, Weissenhofer A, Wanek W (2020) Climatic and edaphic controls over tropical forest diversity and vegetation carbon storage. Sci Rep 10:5066.

Holland N, Richardson AD (2009) Stomatal Length Correlates with Elevation of Growth in Four Temperate Species†. J Sustainable For 28:63–73.

Hubau W, Lewis SL, Phillips OL, Affum-Baffoe K, Beeckman H, Cuní-Sanchez A, Daniels AK, Ewango CEN, Fauset S, Mukinzi JM, Sheil D, Sonké B, Sullivan MJP, Sunderland TCH, Taedoumg H, Thomas SC, White LJT, Abernethy KA, Adu-Bredu S, Amani CA, Baker TR, Banin LF, Baya F, Begne SK, Bennett AC, Benedet F, Bitariho R, Bocko YE, Boeckx P, Boundja P, Brienen RJW, Brncic T, Chezeaux E, Chuyong GB, Clark CJ, Collins M, Comiskey JA, Coomes DA, Dargie GC, de Haulleville T, Kamdem MND, Doucet J-L, Esquivel-Muelbert A, Feldpausch TR, Fofanah A, Foli EG, Gilpin M, Gloor E, Gonmadje C, Gourlet-Fleury S, Hall JS, Hamilton AC, Harris DJ, Hart TB, Hockemba MBN, Hladik A, Ifo SA, Jeffery KJ, Jucker T, Yakusu EK, Kearsley E, Kenfack D, Koch A, Leal ME, Levesley A, Lindsell JA, Lisingo J, Lopez-Gonzalez G, Lovett JC, Makana J-R, Malhi Y, Marshall AR, Martin J, Martin EH, Mbayu FM, Medjibe VP, Mihindou V, Mitchard ETA, Moore S, Munishi PKT, Bengone NN, Ojo L, Ondo FE, Peh KS-H, Pickavance GC, Poulsen AD, Poulsen JR, Qie L, Reitsma J, Rovero F, Swaine MD, Talbot J, Taplin J, Taylor DM, Thomas DW, Toirambe B, Mukendi JT, Tuagben D, Umunay PM, van der Heijden GMF, Verbeeck H, Vleminckx J, Willcock S, Wöll H, Woods JT, Zemagho L (2020) Asynchronous carbon sink saturation in African and Amazonian tropical forests. Nature 579:80–87.

Hultine KR, Marshall JD (2000) Altitude trends in conifer leaf morphology and stable carbon isotope composition. Oecologia 123:32–40.

Hu J-J, Xing Y-W, Turkington R, Jacques FMB, Su T, Huang Y-J, Zhou Z-K (2015) A new positive relationship between pCO_2_ and stomatal frequency in Quercus guyavifolia (Fagaceae): a potential proxy for palaeo-CO2 levels. Ann Bot 115:777–788.

James JC, Grace J, Hoad SP (1994) Growth and Photosynthesis of Pinus Sylvestris at its Altitudinal Limit in Scotland. J Ecol 82:297–306.

Körner C (2007) The use of ‘altitude’ in ecological research. Trends Ecol Evol 22:569–574.

Korner CK, Bannister P, Mark AF (1986) Altitudinal variation in stomatal conductance, nitrogen content and leaf anatomy in different plant life forms in New Zealand. Oecologia 69:577– 588.

Korner C, Cochrane PM (1985) Stomatal responses and water relations of Eucalyptus paucitlora in summer along an elevational gradient. Oecologia 66:443–455.

Kostermans A (1980) Clusiaceae (Guttiferae). In: Dassanayake MD, Fosberg FR (eds) A Revised Handbook to the Flora of Ceylon: Volume I. Amarind, New Delhi, pp 72–110.

Kouwenberg LLR, Kürschner WM, McElwain JC (2007) Stomatal Frequency Change Over Altitudinal Gradients: Prospects for Paleoaltimetry. Rev Mineral Geochem 66:215–241.

Lieberman D, Lieberman M, Peralta R, Hartshorn GS (1996) Tropical Forest Structure and Composition on a Large-Scale Altitudinal Gradient in Costa Rica. J Ecol 84:137–152.

Li C, Zhang X, Liu X, Luukkanen O, Berninger F (2006) Leaf morphological and Pphysiological responses of Quercus aquifolioides along an altitudinal gradient. Silva Fennica 40:5–13.

Lomax BH, Woodward FI, Leitch IJ, Knight CA, Lake JA (2009) Genome size as a predictor of guard cell length in Arabidopsis thaliana is independent of environmental conditions. New Phytol 181:311–314.

Luo J, Zang R, Li C (2006) Physiological and morphological variations of Picea asperata populations originating from different altitudes in the mountains of southwestern China. For Ecol Manage 221:285–290.

Malhi Y, Aragão LEOC, Galbraith D, Huntingford C, Fisher R, Zelazowski P, Sitch S, McSweeney C, Meir P (2009) Exploring the likelihood and mechanism of a climate-change-induced dieback of the Amazon rainforest. Proc Natl Acad Sci USA 106:20610–20615.

Malhi Y, Gardner TA, Goldsmith GR, Silman MR, Zelazowski P (2014) Tropical Forests in the Anthropocene. Annual Review of Environment and Resources 39:125–159.

Malhi Y, Girardin CAJ, Goldsmith GR, Doughty CE, Salinas N, Metcalfe DB, Huaraca Huasco W, Silva-Espejo JE, Del Aguilla-Pasquell J, Farfán Amézquita F, Aragão LEOC, Guerrieri R, Ishida FY, Bahar NHA, Farfan-Rios W, Phillips OL, Meir P, Silman M (2017) The variation of productivity and its allocation along a tropical elevation gradient: a whole carbon budget perspective. New Phytol 214:1019–1032.

Malhi Y, Silman M, Salinas N, Bush M, Meir P, Saatchi S (2010) Introduction: Elevation gradients in the tropics: laboratories for ecosystem ecology and global change research. Glob Chang Biol 16:3171–3175.

McElwain JC (2004) Climate-independent paleoaltimetry using stomatal density in fossil leaves as a proxy for CO_2_ partial pressure. Geology 32:1017–1020.

Meijer W (1983) Anacardiaceae. In: Dassanayake MD, Fosberg FR (eds) A Revised Handbook to the Flora of Ceylon: Volume IV. Amarind, New Delhi

Motsara MR, Roy RN (2008) Guide to Laboratory establishment for plant nutrient analysis. Food and Agriculture Organization, Rome.

Niinemets Ü (2001) Global-scale climatic controls of leaf dry mass per area, density, and thickness in trees and shrubs. Ecology 82:453–469.

Oliveras I, Bentley L, Fyllas NM, Gvozdevaite A, Shenkin AF, Prepah T, Morandi P, Peixoto KS, Boakye M, Adu-Bredu S, Schwantes Marimon B, Marimon Junior BH, Martin R, Asner G, Díaz S, Enquist BJ, Malhi Y (2020) The Influence of Taxonomy and Environment on Leaf Trait Variation Along Tropical Abiotic Gradients. Frontiers in Forests and Global Change 3:18.

Ong CK, Black CR, Saffell RA (1985) Influence of Saturation Deficit on Leaf Production and Expafision in Stands of Groundnut-{Arachis hypogaea L.) Grown Without Irrigation. Ann Bot 56:523–536.

Onoda Y, Wright IJ (2018) The Leaf Economics Spectrum and its Underlying Physiological and Anatomical Principles. In: Adams WW III, Terashima I (eds) The Leaf: A Platform for Performing Photosynthesis. Springer International Publishing, Cham, pp 451–471.

Osnas JLD, Lichstein JW, Reich PB, Pacala SW (2013) Global leaf trait relationships: mass, area, and the leaf economics spectrum. Science 340:741–744.

Qiang W-Y, Wang X-L, Chen T, Feng H-Y, An L-Z, He Y-Q, Wang G (2003) Variations of stomatal density and carbon isotope values of Picea crassifolia at different altitudes in the Qilian Mountains. Trees 17:258–262.

Reich PB, Oleksyn J (2004) Global patterns of plant leaf N and P in relation to temperature and latitude. Proc Natl Acad Sci USA 101:11001–11006.

Salah HBH, Tardieu F (1996) Quantitative analysis of the combined effects of temperature, evaporative demand and light on leaf elongation rate in well-watered field and laboratory-grown maize plants. J Exp Bot 47:1689–1698.

Sanjeewani HKN, Samarasinghe DP, Jayasinghe HD, Gardiyawasam PH, Wahala WMPSB, Wijetunga WMGASTB, Ukuwela KDB, Gomes P, De Costa WAJM (2020) Response of tree community composition, plant diversity and aboveground tree biomass in tropical rainforests of Sri Lanka to variation in altitude. Tropical Agricultural Research 31:87–101.

Shi Z, Haworth M, Feng Q, Cheng R, Centritto M (2015) Growth habit and leaf economics determine gas exchange responses to high elevation in an evergreen tree, a deciduous shrub and a herbaceous annual. AoB Plants 7. http://dx.doi.org/10.1093/aobpla/plv115

Squire GR (1990) The Physiology of Tropical Crop Production. C.A.B. International, Wallingford.

Squire GR, Black CR, Ong CK (1983) Response to saturation deficit of leaf extension in a stand of Pearl Millet (Pennisetum typhoides S. & H.)II: II. Dependence on leaf water status and irradiance. J Exp Bot 34:856–865.

Squire GR, Ong CK (1983) Response to Saturation Deficit of Leaf Extension in a Stand of Pearl Millet (Pennisetum typhoides S. & H.)I: I. Interaction with temperature. J Exp Bot 34:846– 855.

Suhr DD (2005) Principal component analysis vs. exploratory factor analysis (paper 203-30). In: Proceedings of the thirtieth annual SAS® users group international conference. p 30.

Tardieu F, Reymond M, Hamard P, Granier C, Muller B (2000) Spatial distributions of expansion rate, cell division rate and cell size in maize leaves: a synthesis of the effects of soil water status, evaporative demand and temperature. J Exp Bot 51:1505–1514.

Tardieu F, Reymond M, Muller B, Granier C, Simonneau T, Sadok W, Welcker C (2005) Linking physiological and genetic analyses of the control of leaf growth under changing environmental conditions. Aust J Agric Res 56:937–946.

Vazquez G JA, Givnish TJ (1998) Altitudinal gradients in tropical forest composition, structure, and diversity in the Sierra de Manantlán. J Ecol 86:999–1020.

Wong SC, Cowan IR, Farquhar GD (1979) Stomatal conductance correlates with photosynthetic capacity. Nature 282:424–426.

Woodward FI (1987) Stomatal numbers are sensitive to increases in CO_2_ from pre-industrial levels. Nature 327:617–618.

Woodward FI, Lake JA, Quick WP (2002) Stomatal development and CO2: ecological consequences. New Phytol 153:477–484.

Wright IJ, Reich PB, Cornelissen JHC, Falster DS, Groom PK, Hikosaka K, Lee W, Lusk CH, Niinemets Ü, Oleksyn J, Others (2005) Modulation of leaf economic traits and trait relationships by climate. Global Ecology Biogeography 14:411–421.

Wright IJ, Reich PB, Westoby M (2001) Strategy shifts in leaf physiology, structure and nutrient content between species of high-and low-rainfall and high-and low-nutrient habitats. Funct Ecol 15:423–434.

Wright IJ, Reich PB, Westoby M, Ackerly DD, Baruch Z, Bongers F, Cavender-Bares J, Chapin T, Cornelissen JHC, Diemer M, Flexas J, Garnier E, Groom PK, Gulias J, Hikosaka K, Lamont BB, Lee T, Lee W, Lusk C, Midgley JJ, Navas M-L, Niinemets U, Oleksyn J, Osada N, Poorter H, Poot P, Prior L, Pyankov VI, Roumet C, Thomas SC, Tjoelker MG, Veneklaas EJ, Villar R (2004) The worldwide leaf economics spectrum. Nature 428:821–827.

Zelazowski P, Malhi Y, Huntingford C, Sitch S, Fisher JB (2011) Changes in the potential distribution of humid tropical forests on a warmer planet. Philos Trans A Math Phys Eng Sci 369:137–160.

Zhang S-B, Zhou Z-K, Hu H, Xu K, Yan N, Li S-Y (2005) Photosynthetic performances of Quercus pannosa vary with altitude in the Hengduan Mountains, southwest China. For Ecol Manage 212:291–301.

