## supplementary figures S1 - S6 for "Stomatal anatomy, leaf structure and nutrients of tropical rainforest tree species respond to altitude in a coordinated manner in accordance with the leaf economics spectrum"

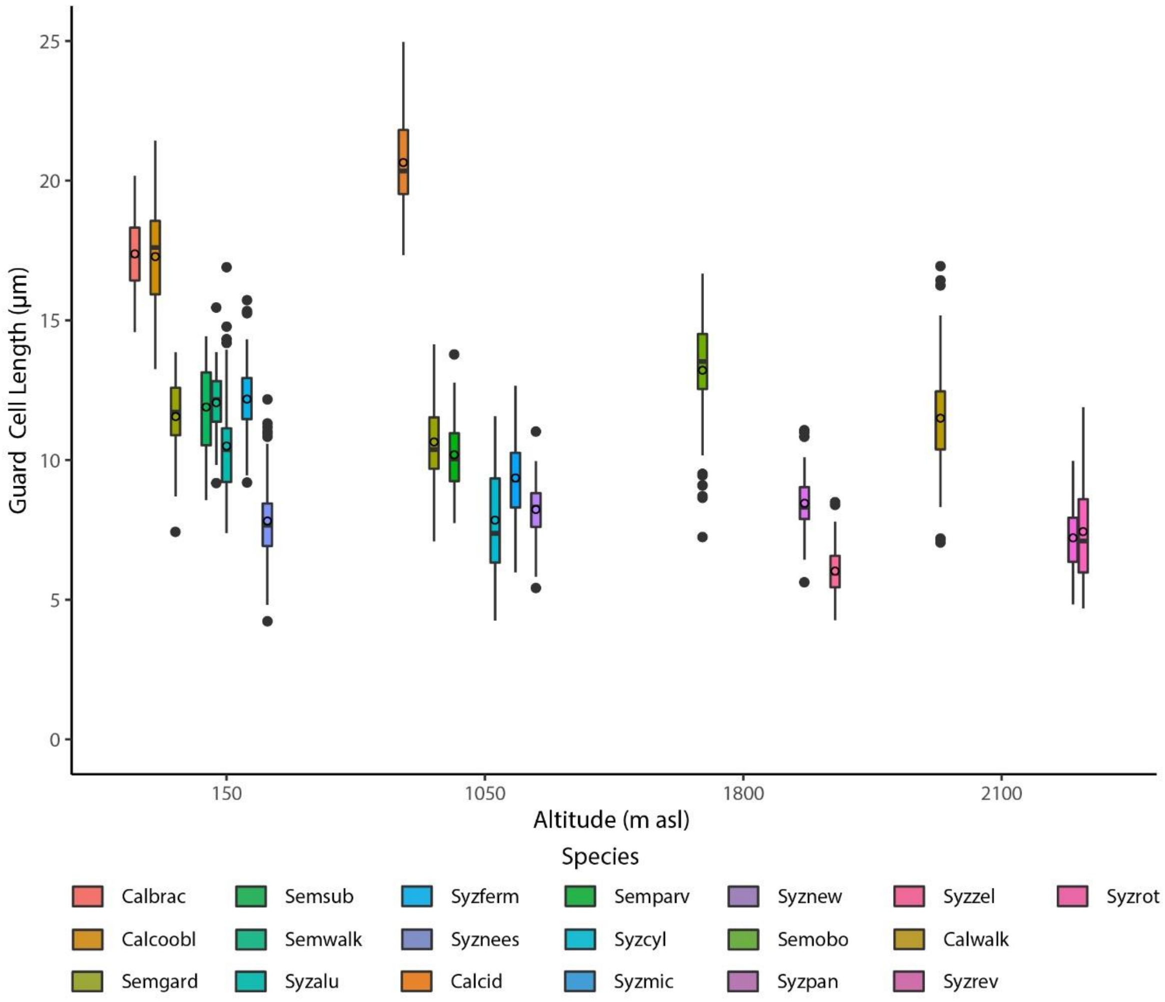


**Figure S1.a** Box plots of distributions of stomatal traits of plant species of tropical rainforests in Sri Lanka at different altitudes : (a) guard cell length; (b) stomatal density; (c) epidermal density; (d) stomatal index; (e) potential conductance index. Abbreviations for species: Calbrac - *Calophyllum bracteatum*; Calcoobl - *Calophyllum cordato-oblongum*; Semgard – *Semecarpus gardneri* (at 150 and 1050 m); Semsub – *Semecarpus subpeltata;* Semwalk – *Semecarpus walkeri*; Syzalu - *Syzygium alubo*; Syzferm - *Syzygium firmum*; Syznees - *Syzygium neesianum*; Calcid - *Calophyllum acidus*; Semparv - *Semecarpus parvifolia*; Syzcyl - *Syzygium cylindricum*; Syzmic - *Syzygium micranthum*; Syznew - *Syzygium spp.*; Semobo - *Semecarpus obovata*; Syzpan - *Syzygium paniculatum*; Syzzel - *Syzygium zeylanicum*; Calwalk - *Calophyllum walkeri*; Syzrev - *Syzygium revolutum*; Syzrot - *Syzygium rotundifolium*.

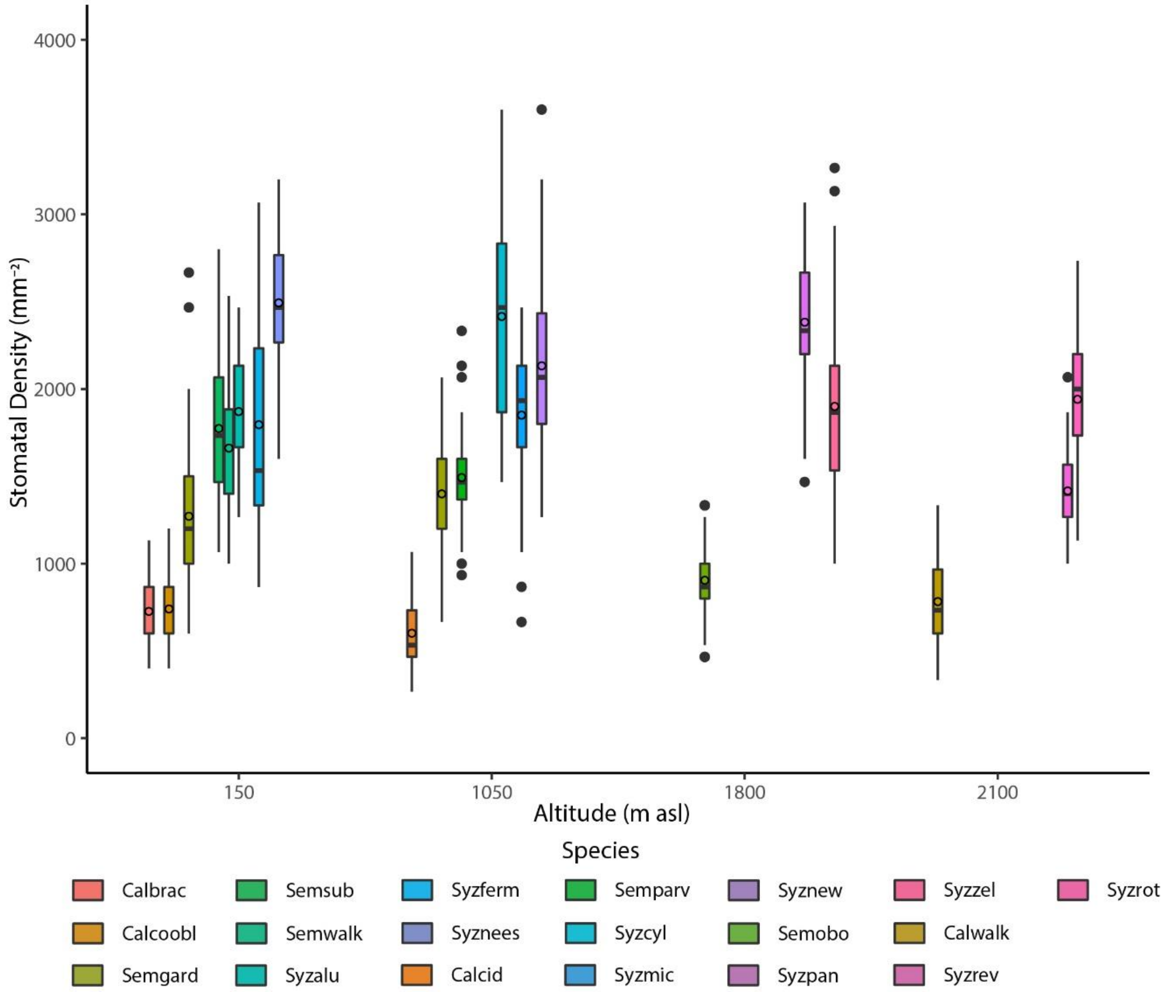


**Figure S1.b** Box plots of distributions of stomatal traits of plant species of tropical rainforests in Sri Lanka at different altitudes : (a) guard cell length; (b) stomatal density; (c) epidermal density; (d) stomatal index; (e) potential conductance index. Abbreviations for species: Calbrac - *Calophyllum bracteatum*; Calcoobl - *Calophyllum cordato-oblongum*; Semgard – *Semecarpus gardneri* (at 150 and 1050 m); Semsub – *Semecarpus subpeltata;* Semwalk – *Semecarpus walkeri*; Syzalu - *Syzygium alubo*; Syzferm - *Syzygium firmum*; Syznees - *Syzygium neesianum*; Calcid - *Calophyllum acidus*; Semparv - *Semecarpus parvifolia*; Syzcyl - *Syzygium cylindricum*; Syzmic - *Syzygium micranthum*; Syznew - *Syzygium spp.*; Semobo - *Semecarpus obovata*; Syzpan - *Syzygium paniculatum*; Syzzel - *Syzygium zeylanicum*; Calwalk - *Calophyllum walkeri*; Syzrev - *Syzygium revolutum*; Syzrot - *Syzygium rotundifolium*.

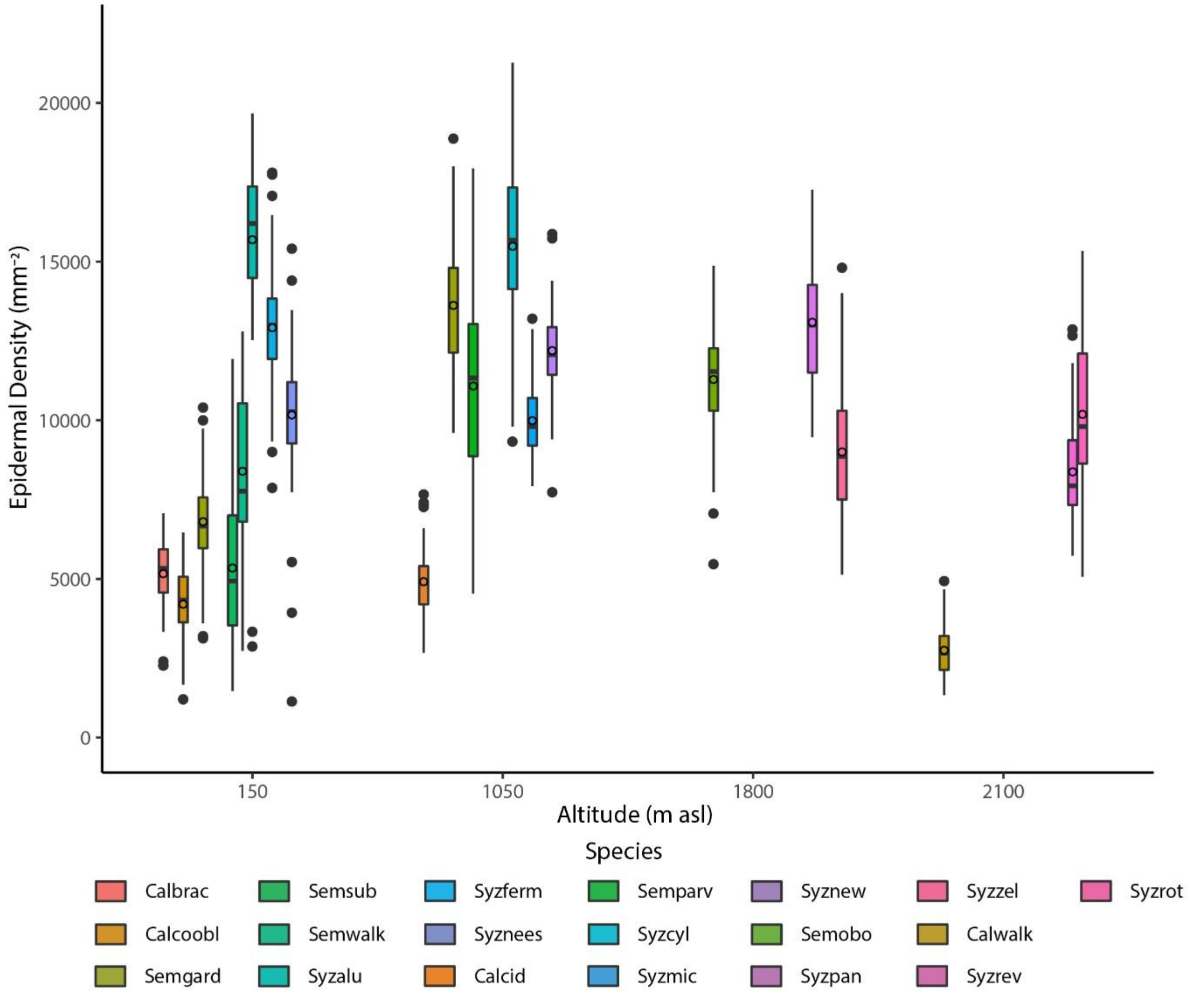


**Figure S1.c** Box plots of distributions of stomatal traits of plant species of tropical rainforests in Sri Lanka at different altitudes : (a) guard cell length; (b) stomatal density; (c) epidermal density; (d) stomatal index; (e) potential conductance index. Abbreviations for species: Calbrac - *Calophyllum bracteatum*; Calcoobl - *Calophyllum cordato-oblongum*; Semgard – *Semecarpus gardneri* (at 150 and 1050 m); Semsub – *Semecarpus subpeltata;* Semwalk – *Semecarpus walkeri*; Syzalu - *Syzygium alubo*; Syzferm - *Syzygium firmum*; Syznees - *Syzygium neesianum*; Calcid - *Calophyllum acidus*; Semparv - *Semecarpus parvifolia*; Syzcyl - *Syzygium cylindricum*; Syzmic - *Syzygium micranthum*; Syznew - *Syzygium spp.*; Semobo - *Semecarpus obovata*; Syzpan - *Syzygium paniculatum*; Syzzel - *Syzygium zeylanicum*; Calwalk - *Calophyllum walkeri*; Syzrev - *Syzygium revolutum*; Syzrot - *Syzygium rotundifolium*.

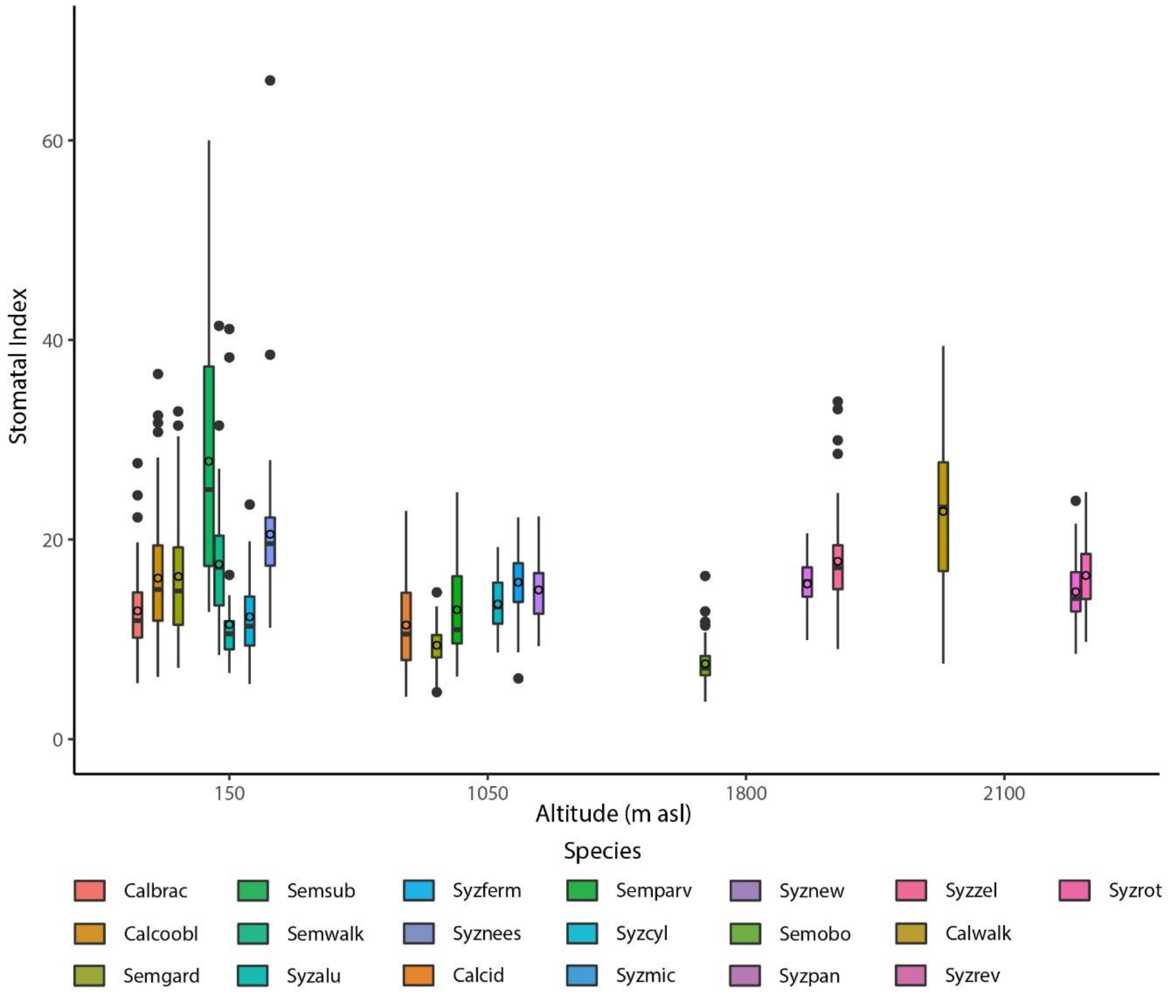


**Figure S1.d** Box plots of distributions of stomatal traits of plant species of tropical rainforests in Sri Lanka at different altitudes : (a) guard cell length; (b) stomatal density; (c) epidermal density; (d) stomatal index; (e) potential conductance index. Abbreviations for species: Calbrac - *Calophyllum bracteatum*; Calcoobl - *Calophyllum cordato-oblongum*; Semgard – *Semecarpus gardneri* (at 150 and 1050 m); Semsub – *Semecarpus subpeltata;* Semwalk – *Semecarpus walkeri*; Syzalu - *Syzygium alubo*; Syzferm - *Syzygium firmum*; Syznees - *Syzygium neesianum*; Calcid - *Calophyllum acidus*; Semparv - *Semecarpus parvifolia*; Syzcyl - *Syzygium cylindricum*; Syzmic - *Syzygium micranthum*; Syznew - *Syzygium spp.*; Semobo - *Semecarpus obovata*; Syzpan - *Syzygium paniculatum*; Syzzel - *Syzygium zeylanicum*; Calwalk - *Calophyllum walkeri*; Syzrev - *Syzygium revolutum*; Syzrot - *Syzygium rotundifolium*.

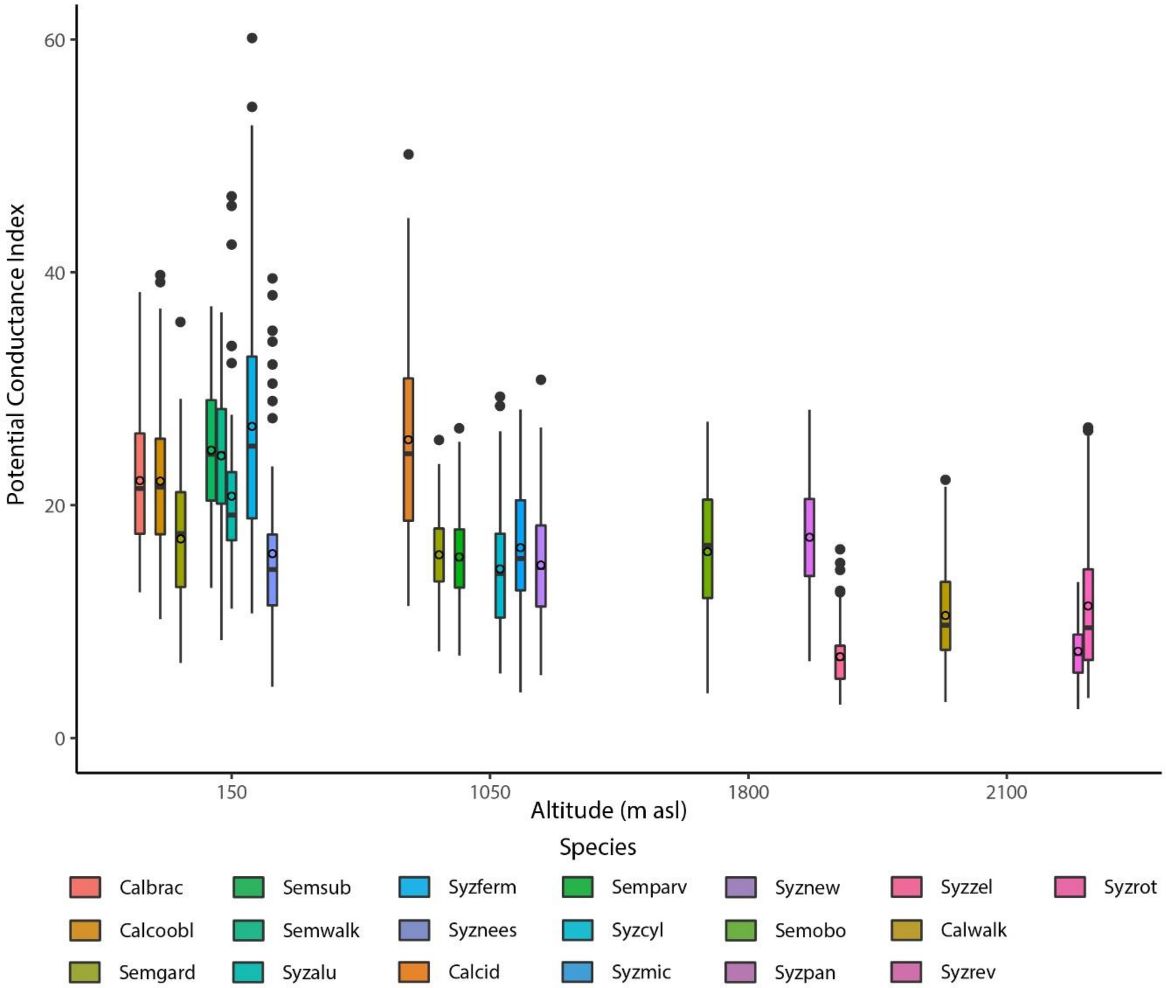


**Figure S1.e** Box plots of distributions of stomatal traits of plant species of tropical rainforests in Sri Lanka at different altitudes : (a) guard cell length; (b) stomatal density; (c) epidermal density; (d) stomatal index; (e) potential conductance index. Abbreviations for species: Calbrac - *Calophyllum bracteatum*; Calcoobl - *Calophyllum cordato-oblongum*; Semgard – *Semecarpus gardneri* (at 150 and 1050 m); Semsub – *Semecarpus subpeltata;* Semwalk – *Semecarpus walkeri*; Syzalu - *Syzygium alubo*; Syzferm - *Syzygium firmum*; Syznees - *Syzygium neesianum*; Calcid - *Calophyllum acidus*; Semparv - *Semecarpus parvifolia*; Syzcyl - *Syzygium cylindricum*; Syzmic - *Syzygium micranthum*; Syznew - *Syzygium spp.*; Semobo - *Semecarpus obovata*; Syzpan - *Syzygium paniculatum*; Syzzel - *Syzygium zeylanicum*; Calwalk - *Calophyllum walkeri*; Syzrev - *Syzygium revolutum*; Syzrot - *Syzygium rotundifolium*.

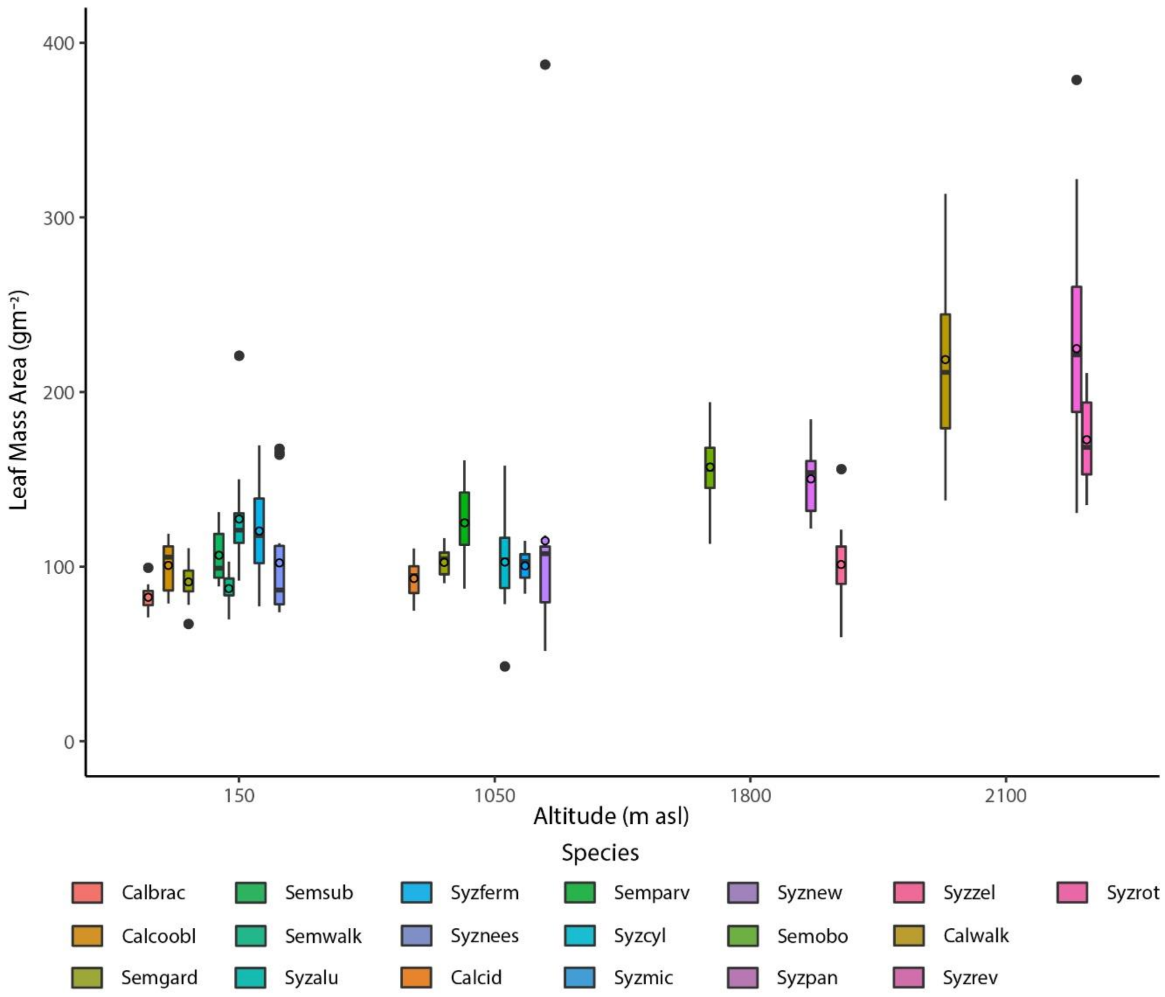


**Figure S2.a** Box plots of distributions of leaf structural traits of plant species of tropical rainforests in Sri Lanka at different altitudes: (a) leaf mass per area; (b) leaf blade area. Abbreviations for species are as in Fig. S.1.

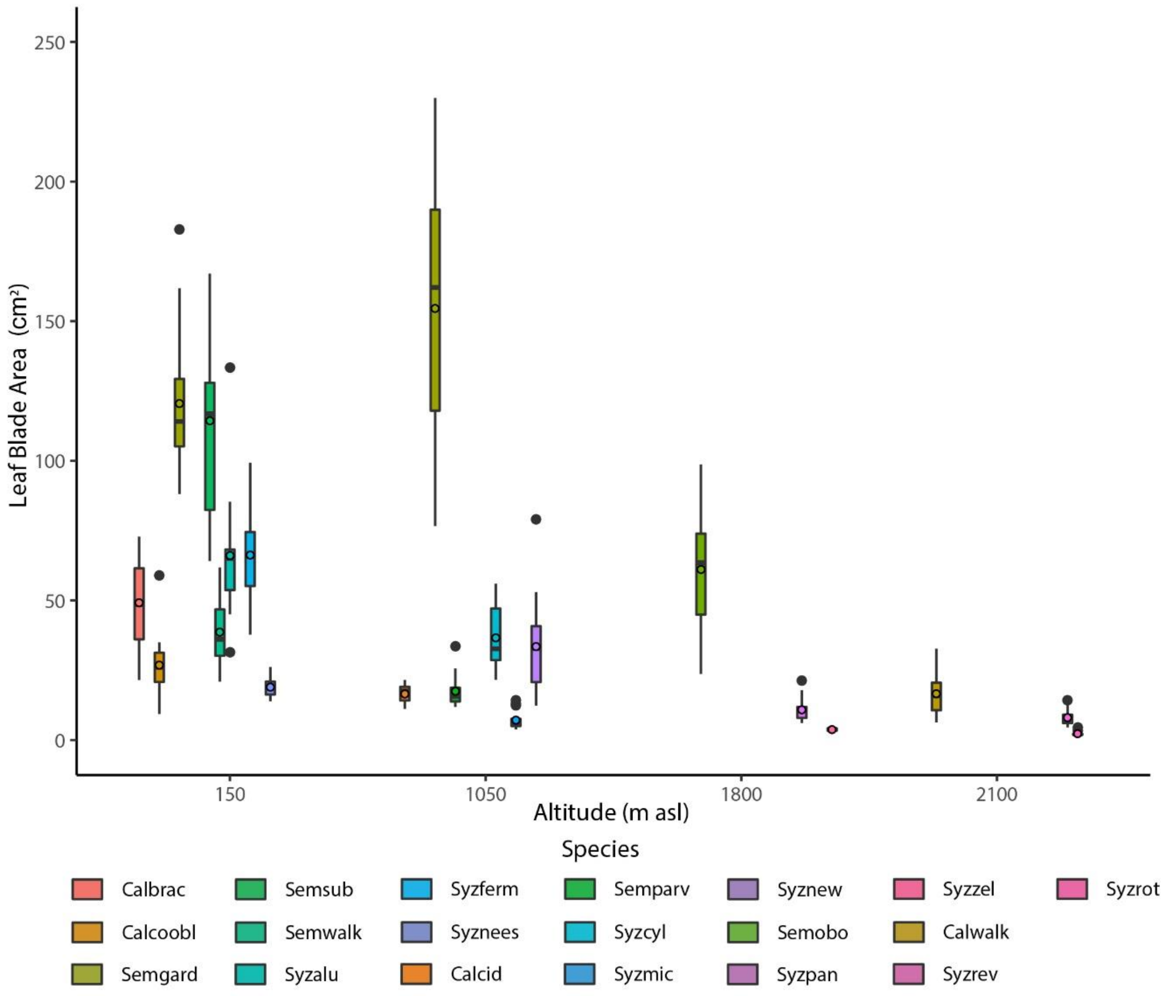


**Figure S2.b** Box plots of distributions of leaf structural traits of plant species of tropical rainforests in Sri Lanka at different altitudes: (a) leaf mass per area; (b) leaf blade area. Abbreviations for species are as in Fig. S.1.

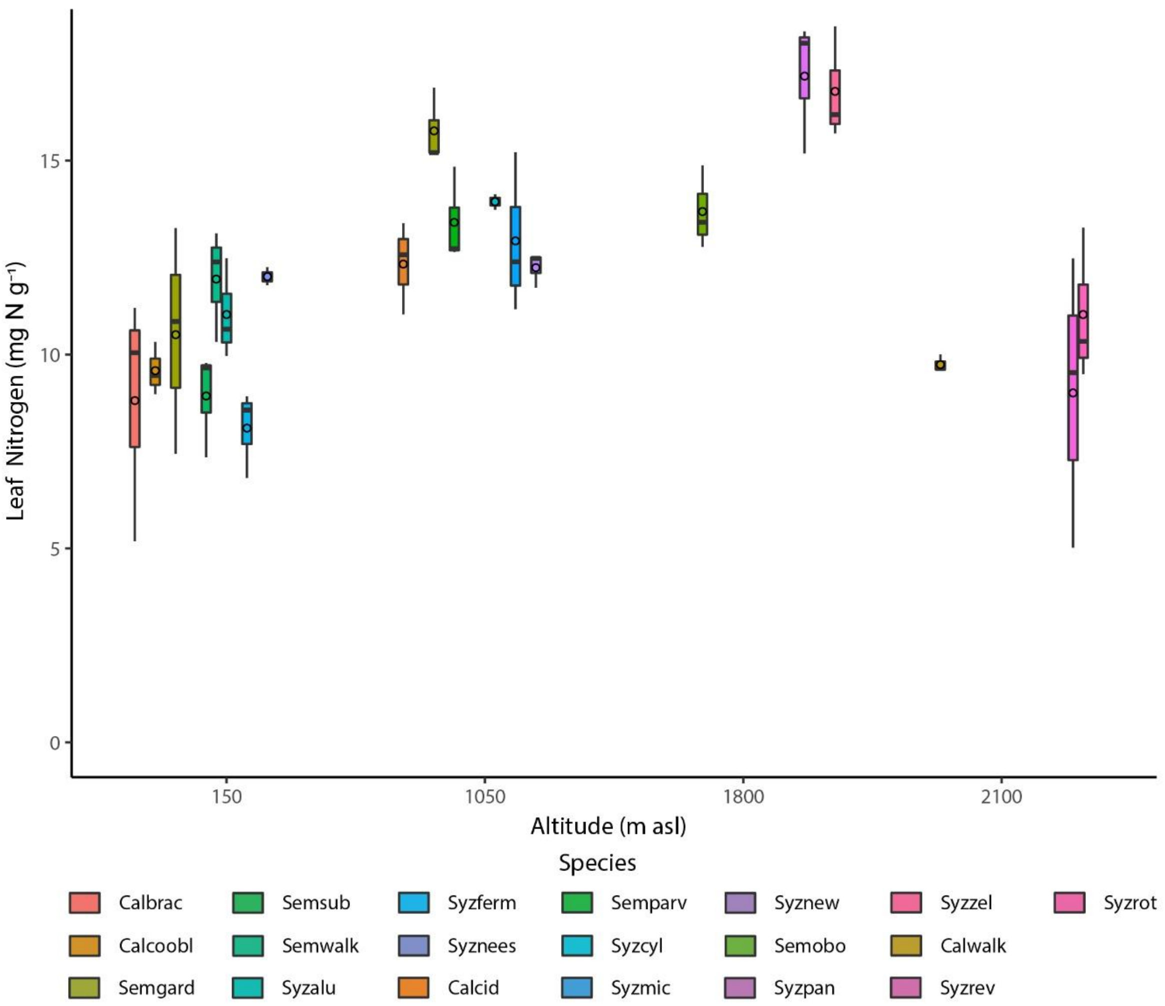


**Figure S3.a** Box plots of distributions of leaf nutrients of plant species of tropical rainforests in Sri Lanka at different altitudes: (a) leaf nitrogen; (b) leaf phosphorus; (c) leaf nitrogen:phosphorus ratio. Abbreviations for species are as in Fig. S.1.

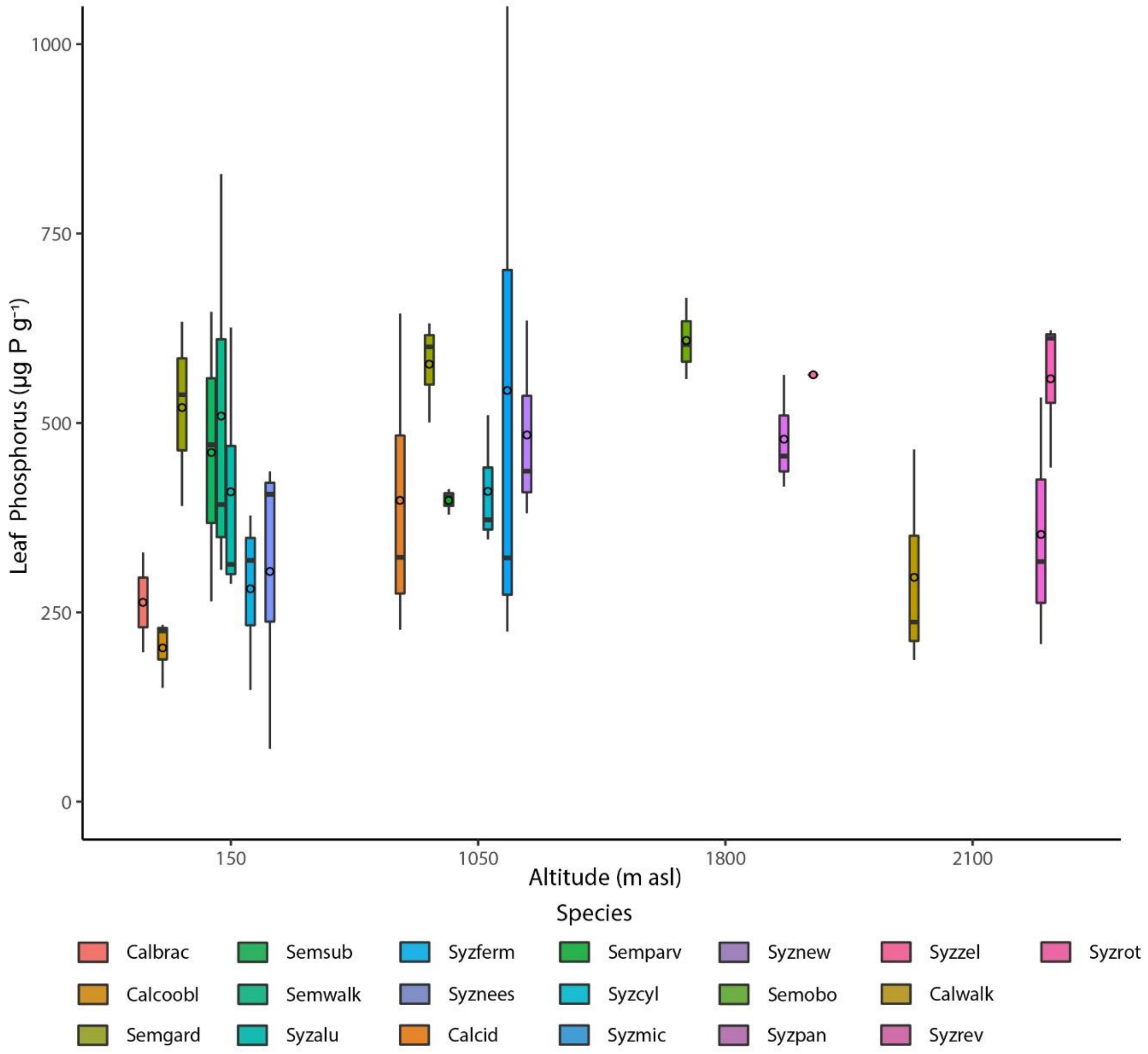


**Figure S3.b** Box plots of distributions of leaf nutrients of plant species of tropical rainforests in Sri Lanka at different altitudes: (a) leaf nitrogen; (b) leaf phosphorus; (c) leaf nitrogen:phosphorus ratio. Abbreviations for species are as in Fig. S.1.

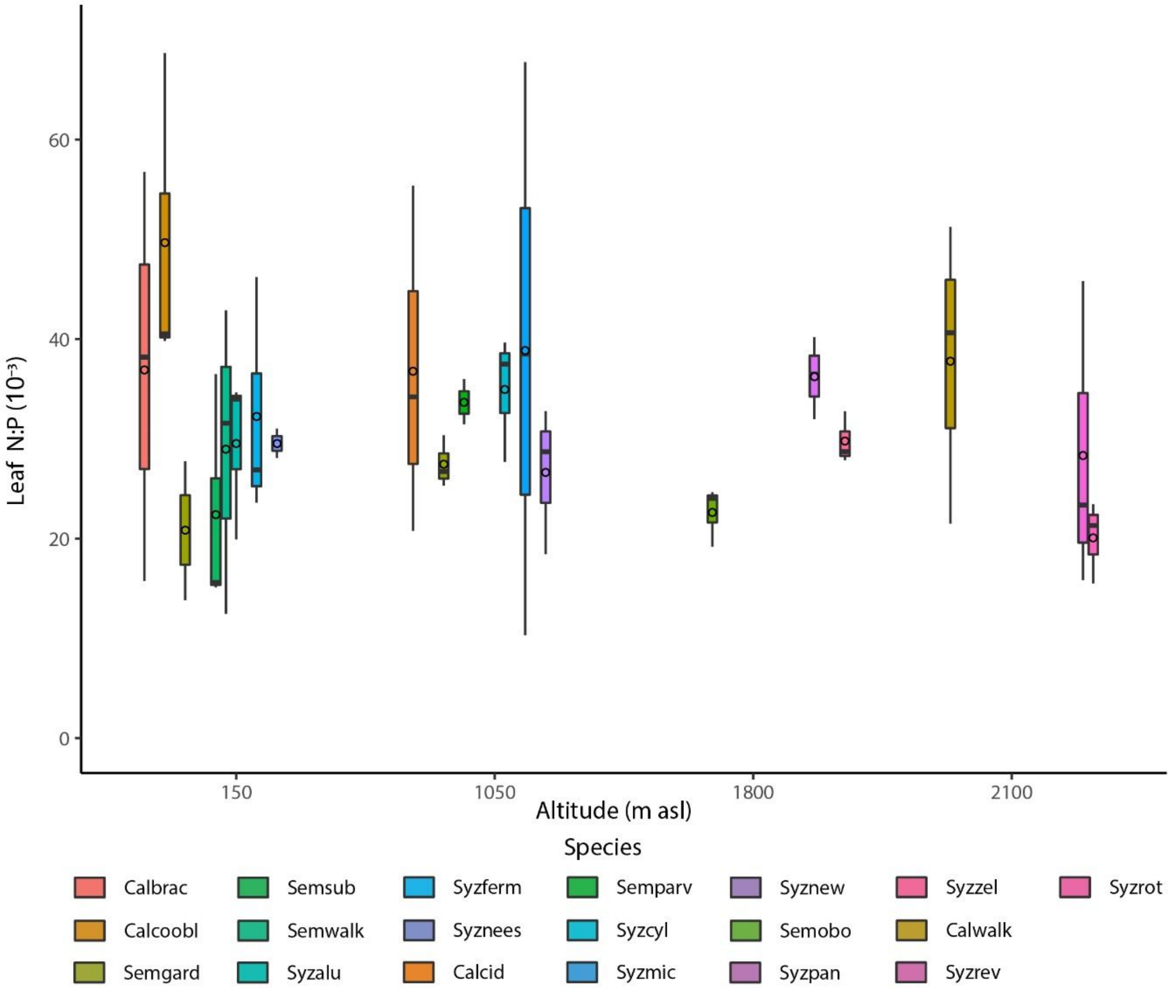


**Figure S3.c** Box plots of distributions of leaf nutrients of plant species of tropical rainforests in Sri Lanka at different altitudes: (a) leaf nitrogen; (b) leaf phosphorus; (c) leaf nitrogen:phosphorus ratio. Abbreviations for species are as in Fig. S.1.

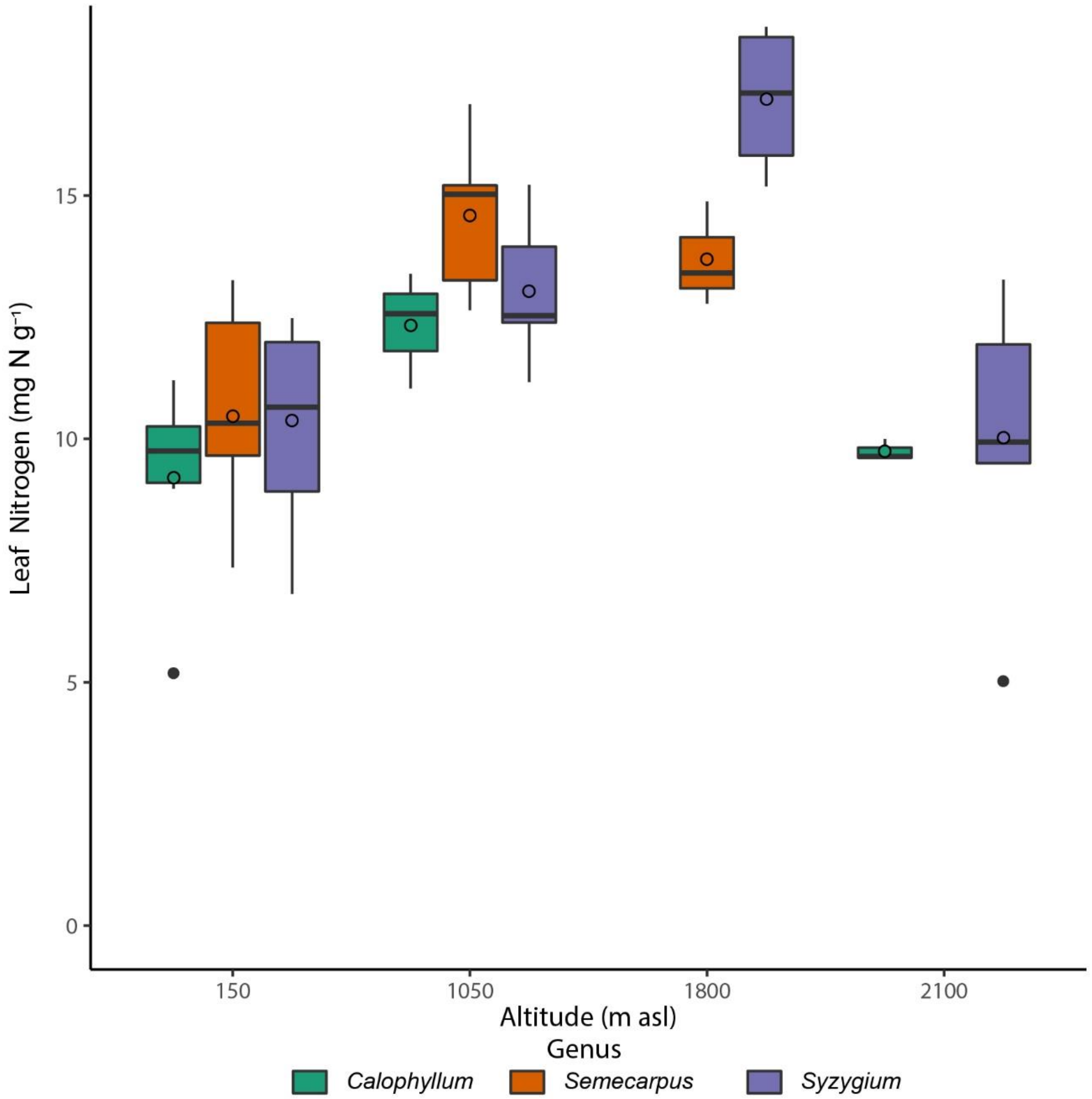


**Figure S4.a** Box plots of distributions of leaf nutrients of plant genera of tropical rainforests in Sri Lanka at different altitudes: (a) leaf nitrogen; (b) leaf phosphorus; (c) leaf nitrogen:phosphorus ratio.

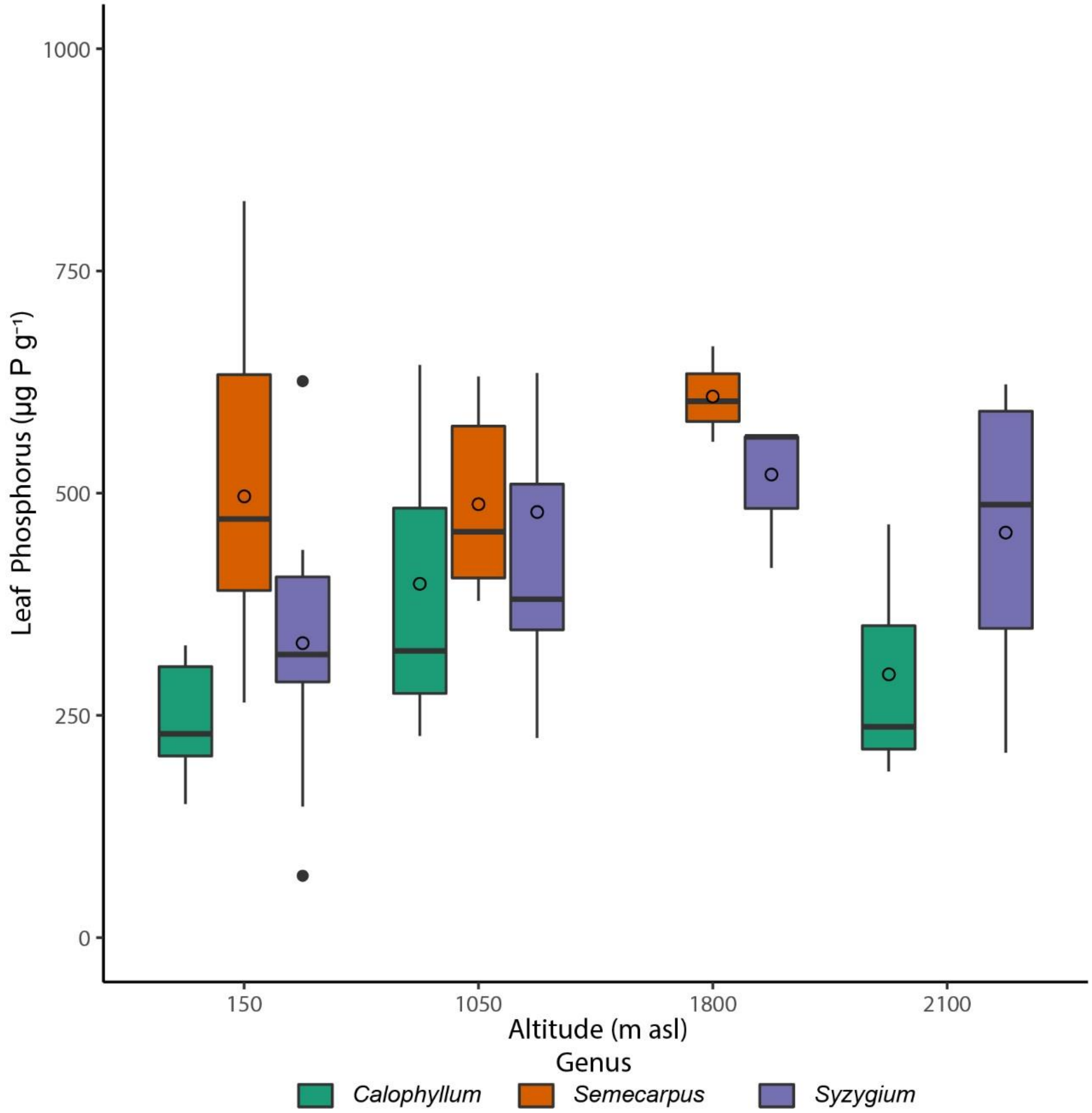


**Figure S4.b** Box plots of distributions of leaf nutrients of plant genera of tropical rainforests in Sri Lanka at different altitudes: (a) leaf nitrogen; (b) leaf phosphorus; (c) leaf nitrogen:phosphorus ratio.

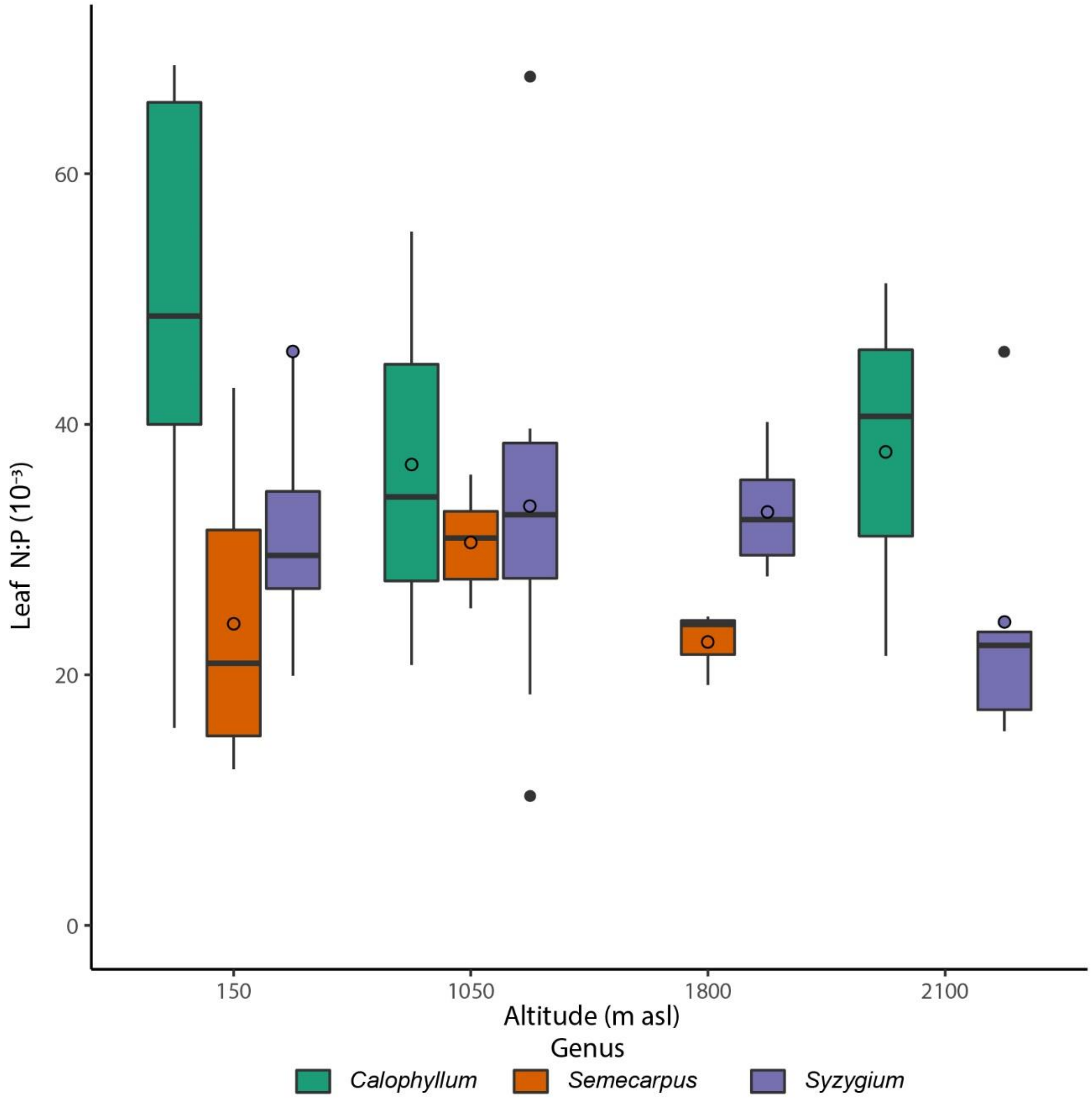


**Figure S4.c** Box plots of distributions of leaf nutrients of plant genera of tropical rainforests in Sri Lanka at different altitudes: (a) leaf nitrogen; (b) leaf phosphorus; (c) leaf nitrogen:phosphorus ratio.

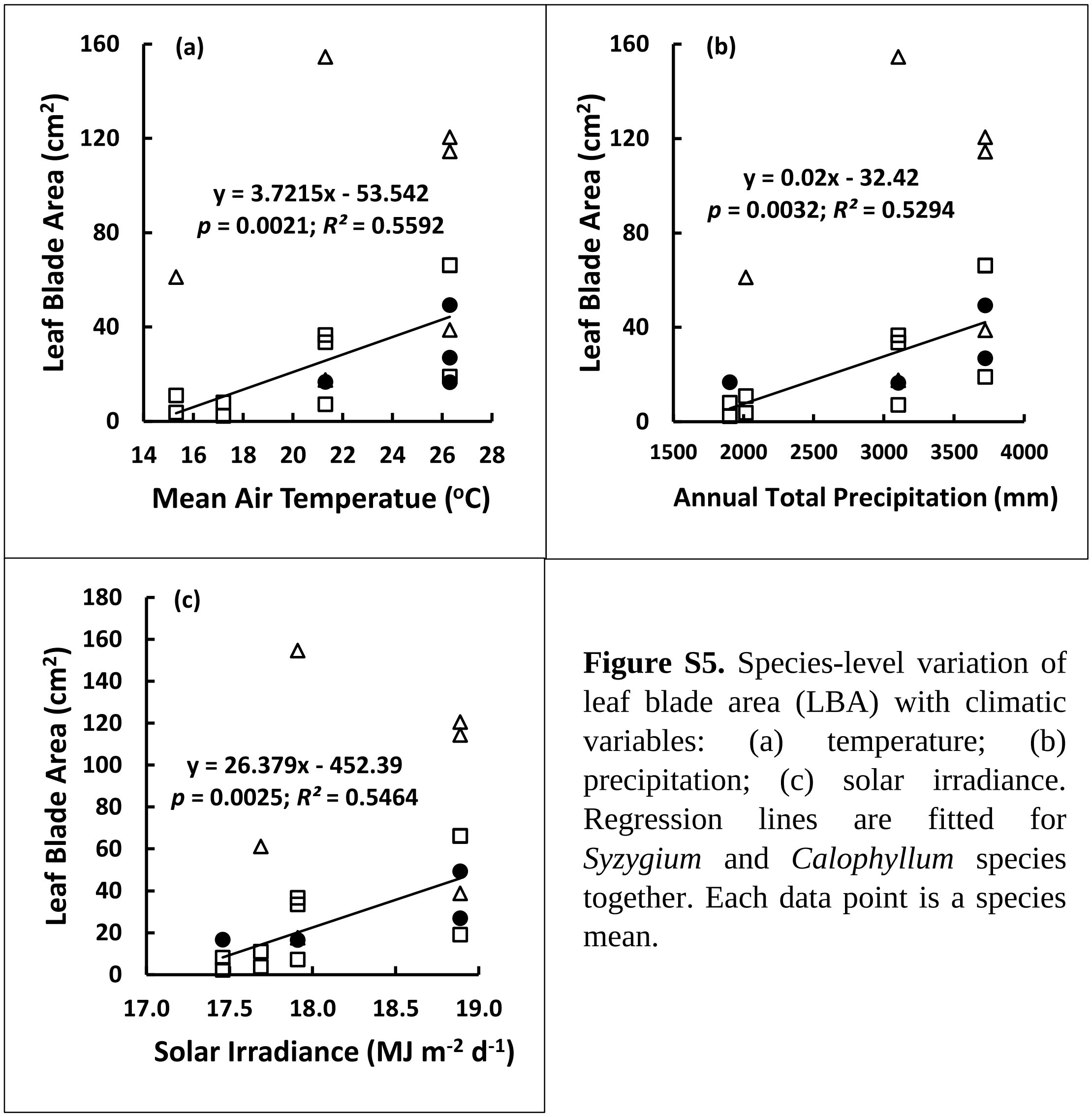


**Figure S5.** Species-level variation of leaf blade area (LBA) with climatic variables: (a) temperature; (b) precipitation; (c) solar irradiance. Regression lines are fitted for *Syzygium* and *Calophyllum* species together. Each data point is a species mean.

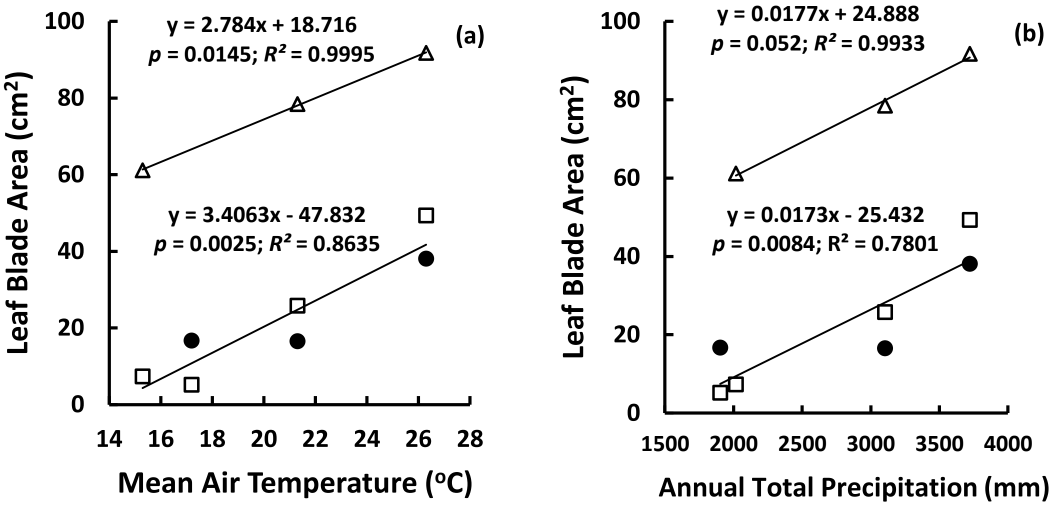


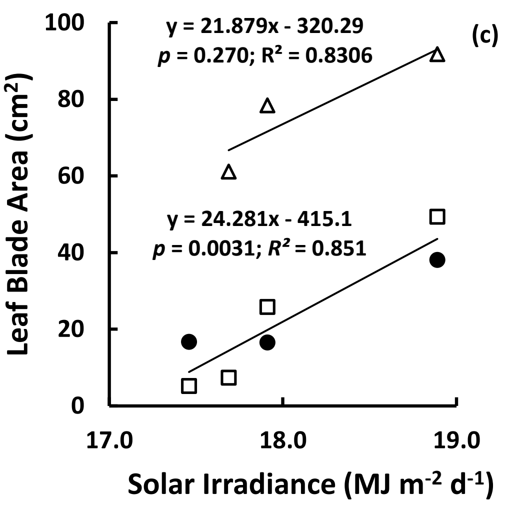


**Figure S6.** Genus-level variation of leaf blade area (LBA) with climatic variables: (a) temperature; (b) precipitation; (c) solar irradiance. Regression lines are fitted for *Semecarpus* separately and for *Syzygium* and *Calophyllum* together. Each data point is the mean of different species of a given genus.
